# Longitudinal miRNA-mRNA Profiling Reveals Network Reorganization in Preeclampsia

**DOI:** 10.64898/2026.11.09.765381

**Authors:** Hooman Mirzakhani, Catherine Jung, Zheng Lu, Diane E. Handy, Joseph Loscalzo, Scott T. Weiss

## Abstract

**Background:** Preeclampsia (PE) is a leading cause of maternal and perinatal morbidity. Temporal dynamics of molecular dysregulation across pregnancy remain poorly understood. We performed longitudinal RNA sequencing of maternal whole blood to characterize miRNA–mRNA regulatory network evolution in PE.

**Methods:** Whole-blood samples from early (M1: 10–18 weeks) and late pregnancy (M2: 32–38 weeks) were analyzed from 43 PE cases and 106 normotensive controls, including 109 paired samples (36 PE cases, 73 controls). Differential expression, temporal classification, pathway enrichment, and miRNA–mRNA network integration were performed. Placental expression support was assessed using internal RNA-seq and public databases. Key miRNA–mRNA interactions were experimentally validated in HTR-8/SVneo trophoblast cells.

**Results:** PE exhibited marked temporal reorganization of miRNA–mRNA networks, with strongly opposing temporal effect sizes across overlapping miRNA and mRNA features (Spearman ρ = −0.82, P < 0.0001). PI3K/AKT/mTORC1 signaling showed biphasic reversal (NES = +1.46, FDR = 0.028 at M1; NES = −1.64, FDR = 0.025 at M2), and heme-metabolism programs showed progressive suppression (2.1-fold NES intensification; FDR < 0.05 at both timepoints). Network analysis identified 29 high-confidence biphasic miRNA–mRNA pairs, including convergent targeting of ADGRA2 by three miRNAs and HSD17B1 by three miRNAs. miR-101-3p overexpression reduced DEPP1 mRNA (∼40%) and protein (∼26%) in trophoblast cells (P < 0.01), supporting a FOXO–DEPP1 axis. Blood-derived pairs showed pathway convergence with placental VGLL3-centered programs (PI3K/AKT, FOXO, TGF-β, trophoblast differentiation).

**Conclusions:** PE is associated with predominantly stage-specific or biphasic miRNA–mRNA network reorganization, with PI3K/AKT/mTORC1 dynamic remodeling and progressive heme-metabolism suppression. These findings support gestational-age-specific biomarker development and identify regulatory axes, including miR-101-3p → DEPP1, for mechanistic investigation.

**Trial registration:** This study is an ancillary analysis from the VDAART, which is registered with ClinicalTrials.gov (NCT00920621: https://clinicaltrials.gov/study/NCT00920621?term=vdaart&rank=1).

## Introduction

Preeclampsia (PE), a hypertensive disorder affecting 2–8% of pregnancies worldwide,^1,2^ remains a major cause of maternal and perinatal morbidity and mortality.^1,3^ It is characterized by new-onset hypertension after 20 weeks of gestation with proteinuria and/or maternal organ dysfunction,^4^ and delivery remains the only definitive treatment.^5^ Improved understanding of PE-associated molecular mechanisms is therefore essential for earlier risk stratification and identification of biologically relevant targets.

Abnormal placentation is central to PE pathophysiology,^6,7^ yet the molecular programs linking early placental dysfunction to later maternal disease remain incompletely understood.^8,9^ Although PE is diagnosed after clinical symptoms emerge, pathogenic processes are thought to begin earlier during trophoblast invasion and spiral artery remodeling in the first half of gestation.^7,10^ A major limitation in the field is the lack of longitudinal molecular profiling across pregnancy. Whether early molecular abnormalities persist, reverse direction, or represent stage-specific responses remains unclear, despite direct relevance to biomarker development and disease interpretation.^11,12^

MicroRNAs (miRNAs) regulate gene expression through post-transcriptional repression of target mRNAs.^13,14^ Because individual miRNAs can regulate multiple transcripts within shared pathways, ^15^ they may shape coordinated regulatory programs rather than isolated single-gene effects. In maternal whole blood, integrated miRNA–mRNA profiling may capture systemic signatures reflecting immune, vascular, and placental-related perturbations during pregnancy.^16,17^ Paired miRNA–mRNA analysis therefore provides a systems-level approach for identifying regulatory circuits associated with PE development and progression.^18,19^

In our previous microarray-based study, we identified early-pregnancy circulating miRNA signatures associated with PE and maternal vitamin D insufficiency.^20^ However, whether PE signatures persist, reverse, or change across pregnancy—and which regulatory programs represent stage-dependent rather than static disease-associated mechanisms—remains unknown. In the present study, we performed miRNA and mRNA sequencing of maternal whole-blood samples collected in early pregnancy, 10–18 weeks, and late pregnancy, 32–38 weeks, from women with and without PE. We identified differentially expressed miRNAs and mRNAs at each time point, evaluated temporal patterns across pregnancy, integrated dysregulated miRNA– mRNA pairs into regulatory networks, and characterized associated biological pathways. We further assessed placental expression of selected blood-derived signatures and experimentally validated the miR-101-3p–DEPP1 regulatory axis in trophoblast cells.

## METHODS

### Study Design and Analytical Overview

**Figure 1** summarizes the study design and analytical workflow. We performed RNA sequencing of miRNA and mRNA from maternal whole blood collected during early pregnancy, 10–18 weeks’ gestation, and late pregnancy, 32–38 weeks’ gestation, from women who subsequently developed preeclampsia and normotensive controls. Differential expression analyses were performed separately at each timepoint, followed by temporal classification of early pregnancy signatures, miRNA–mRNA target integration, and pathway enrichment analysis. We further assessed placental expression support for selected blood-derived signatures and experimentally evaluated a prioritized miRNA–mRNA interaction, miR-101-3p → *DEPP1*, in trophoblast cells. Expanded protocol details are provided in the **Supplementary Methods**.

**Figure 1.**
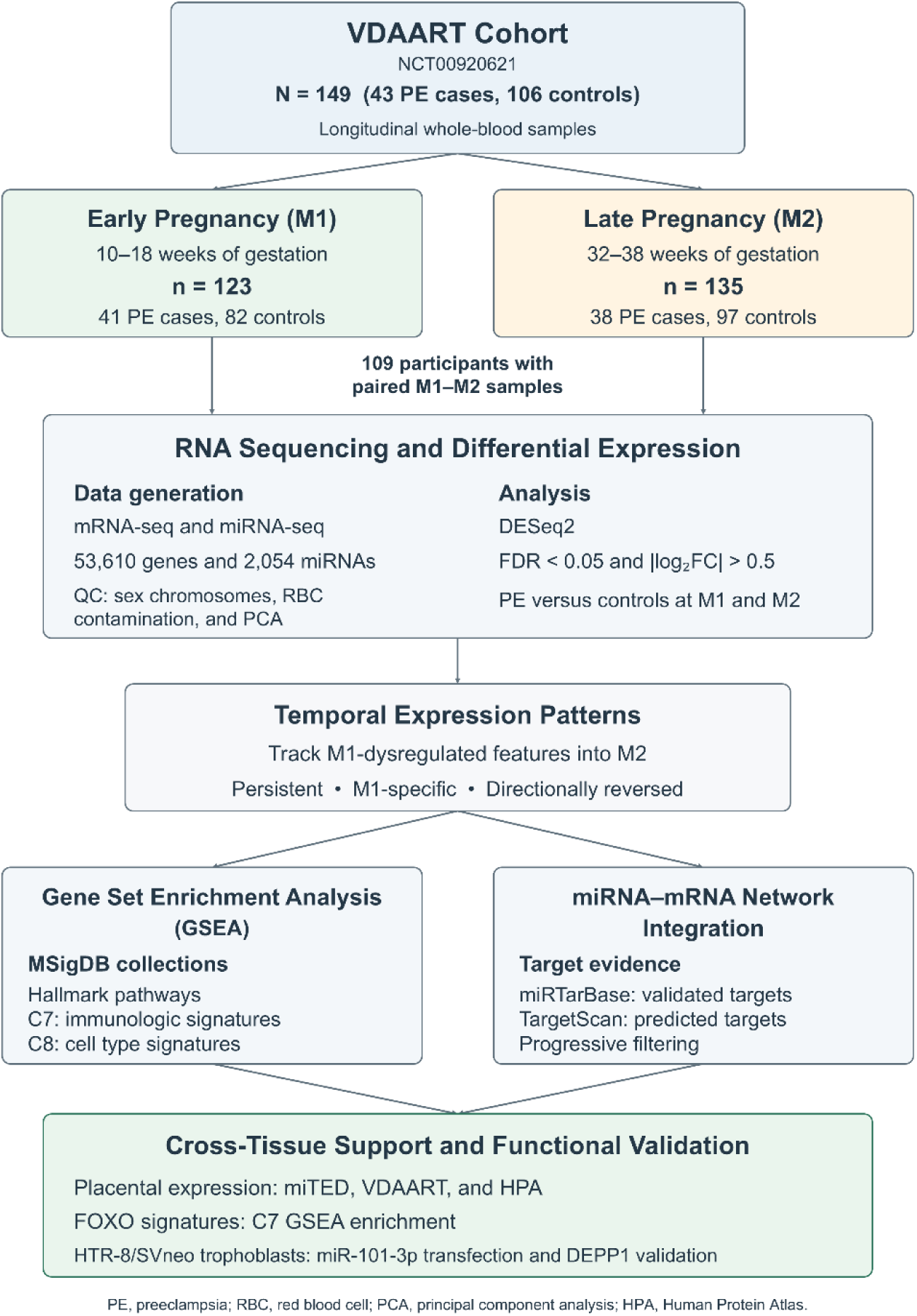
Study design and analytical workflow. Longitudinal whole blood samples were collected from the VDAART cohort (NCT00920621) at early pregnancy (M1: 10–18 weeks, asymptomatic) and late pregnancy (M2: 32–38 weeks, symptomatic), with 109 participants providing paired samples. RNA sequencing profiled mRNA and miRNA expression (53,610 genes, 2,054 miRNAs), followed by quality control and differential expression analysis (DESeq2; FDR < 0.05, |log₂FC| > 0.5). Temporal dynamics analysis tracked M1-dysregulated features into M2. Gene set enrichment analysis (GSEA) interrogated MSigDB Hallmark, immunologic (C7), and cell type (C8) signatures. miRNA-target network integration used miRTarBase and TargetScan databases. Multi-level validation included placental expression profiling (miTED, VDAART, HPA), FOXO pathway signatures, and experimental confirmation via miR-101-3p transfection in HTR-8/SVneo cells. PE, preeclampsia; QC, quality control; chr, chromosome; RBC, red blood cell; PCA, principal component analysis; HPA, Human Protein Atlas.

### Parent Cohort and Study Participants

The Vitamin D Antenatal Asthma Reduction Trial (VDAART; NCT00920621) was a multicenter, randomized, placebo-controlled trial sponsored by the National Heart, Lung, and Blood Institute. Pregnant participants with singleton pregnancies were enrolled between 10 and 18 weeks’ gestation and randomized to receive either 4,400 IU/day vitamin D3, consisting of 4,000 IU/day cholecalciferol plus 400 IU/day prenatal vitamin, or 400 IU/day prenatal vitamin alone. The trial was approved by the Institutional Review Boards at all participating clinical centers and the Data Coordinating Center at Brigham and Women’s Hospital, and all participants provided written informed consent. Additional trial design details and trial outcomes have been reported previously.^21–23^

Preeclampsia was the secondary outcome of the trial, and the present study used a nested case-control design within VDAART.^24^ Among participants who developed PE, those with available maternal whole-blood samples from early pregnancy were eligible. Controls were selected from VDAART participants without pregnancy complications and matched to cases on maternal age, self-reported race, enrollment site, and treatment arm. Additional late-pregnancy samples were obtained when available to support longitudinal analysis. After RNA extraction, sequencing, and quality-control filtering, 43 PE cases and 106 controls had usable samples at one or both timepoints, including 41 PE cases and 82 controls at M1, 38 PE cases and 97 controls at M2, and 36 PE cases and 73 controls with paired M1 and M2 samples.

### Preeclampsia Definition and Clinical Characterization

PE was a prespecified secondary outcome and adverse event in VDAART. After delivery, medical records were systematically abstracted, and potential cases were reviewed by a blinded adjudication committee of board-certified obstetricians without knowledge of treatment assignment.^24^ PE was defined according to American College of Obstetricians and Gynecologists (ACOG) criteria as new-onset hypertension after 20 weeks’ gestation with proteinuria and/or evidence of maternal organ dysfunction. Severe PE was defined by the presence of severe features, including severe-range blood pressure, thrombocytopenia, elevated liver enzymes, renal insufficiency, pulmonary edema, or cerebral/visual symptoms.^4^ Gestational age at delivery and preterm birth status were abstracted from medical records. Maternal 25-hydroxyvitamin D [25(OH)D] concentrations were measured at enrollment and late pregnancy and were used in descriptive and sensitivity analyses.

### Whole-Blood RNA Extraction, Library Preparation, and Sequencing

Maternal whole blood was collected in PAXgene Blood RNA tubes. Total RNA, including small RNAs, was extracted using the PAXgene Blood miRNA Kit. RNA quantity and quality were assessed before library preparation, and samples meeting prespecified thresholds for RNA integrity and concentration were advanced to sequencing.

For mRNA-seq, stranded total RNA libraries were prepared using ribosomal RNA and globin depletion and sequenced on the Illumina NextSeq 550 platform to generate paired-end 75-bp reads, targeting approximately 75 million reads per sample. For miRNA-seq, small RNA libraries were prepared using the NEXTflex Small RNA Sequencing Kit and sequenced on the Illumina NextSeq 550 platform to generate single-end 75-bp reads, targeting approximately 12 million reads per sample. Erythrocyte miRNA blockers (miR-486-5p, has-miR-92a, and has-miR-451a) were used during small RNA library preparation to reduce contamination from highly abundant hemolysis-associated miRNAs.^25^ Detailed library preparation, sequencing, and quality-control procedures are provided in the **Supplementary Methods**.

### Sequencing Data Processing and Feature Filtering

Raw mRNA-seq and miRNA-seq reads were processed using bcbio-nextgen.^26^ For mRNA-seq, reads were quality assessed, adapter trimmed, aligned to the human reference genome GRCh38 using STAR, and quantified at the gene level using featureCounts with GENCODE v46 annotations. For miRNA-seq, reads were quality assessed, adapter trimmed, filtered for mature miRNA-compatible read lengths, aligned to miRBase release 22, and quantified for known mature miRNAs. Samples were excluded for low alignment rate or insufficient sequencing depth according to prespecified thresholds.

Count matrices were imported into R for downstream analysis. Genes and miRNAs located on chromosomes X and Y were excluded to reduce potential confounding by sex chromosome dosage and X-inactivation. Features were retained if they had at least 1 count in at least 50% of samples within comparison groups. Variance-stabilizing transformation was used for exploratory analyses and visualization, including principal component analysis.

### Differential Expression Analysis

Differential expression analyses were performed using DESeq2.^27^ Analyses were conducted separately for early pregnancy, M1, and late pregnancy, M2. At each timepoint, PE case-control status was modeled as the primary predictor. Primary differential expression models adjusted for fetal sex, maternal race/ethnicity, treatment arm, gestational age at sampling, sequencing batch, maternal asthma, body mass index (BMI) at enrollment, and parity. PE cases had lower early-pregnancy 25-hydroxyvitamin D [25(OH)D] concentrations and higher body mass index at enrollment. Body mass index was included as a covariate in the primary differential expression models, while 25(OH)D was evaluated in sensitivity analyses because of its baseline difference between groups and its inverse correlation with BMI. Differentially expressed genes and miRNAs were defined as features with Benjamini–Hochberg false discovery rate (FDR) <0.05 and absolute log2 fold change >0.5.

Although differential expression testing was performed for all expressed annotated features, including protein-coding genes, long noncoding RNAs, pseudogenes, and other transcripts, downstream pathway enrichment and miRNA–mRNA integration focused on protein-coding genes to support pathway interpretation and target-based regulatory analysis.

### Temporal Classification of Early Pregnancy Signatures

To distinguish stable from stage-dependent molecular signatures, we identified features differentially expressed at M1 and evaluated their corresponding behavior at M2. M1-dysregulated features were classified as same-direction persistent if they also met significance criteria at M2 in the same direction, biphasic if they also met significance criteria at M2 in the opposite direction, or M1-specific if they did not meet significance criteria at M2. Temporal classification was performed for protein-coding genes, miRNAs, and the full set of M1-dysregulated annotated features. Correlation of effect sizes between M1 and M2 was evaluated using Spearman correlation.

### Red Blood Cell Contamination and Hemolysis Assessment

Because heme-metabolism pathways were identified in downstream analyses, we evaluated whether heme-related findings could reflect differential erythroid contamination or hemolysis. We calculated single-sample enrichment scores for a 20-gene erythroid marker signature and a 9-gene hemoglobin signature using variance-stabilized expression values.^28^ Associations between each signature and PE status were tested separately at M1 and M2 using logistic regression models adjusted for clinical covariates. Detailed gene lists and modeling procedures are provided in **Supplementary Appendix** and **Supplementary Tables: Tables S2 and S3**.

### Gene Set Enrichment Analysis

Pathway enrichment analysis was performed on ranked protein-coding genes using clusterProfiler.^29^ Curated gene sets were obtained from the Molecular Signatures Database, including Hallmark, Immunologic Signatures, and Cell Type Signatures.^30^ Preranked GSEA was performed separately at M1 and M2 by ranking expressed protein-coding genes according to the differential expression test statistic. Gene sets were considered significantly enriched at FDR <0.05, and normalized enrichment scores were used to determine direction and magnitude of enrichment. Enriched Hallmark, C7, and C8 GSEA results are provided in **Supplementary Data 2**.

### miRNA–mRNA Target Prediction and Network Integration

Putative miRNA–mRNA regulatory relationships were identified using multiMiR, which integrates experimentally validated and computationally predicted miRNA-target databases.^31^ Candidate pairs were retained when a differentially expressed miRNA had a predicted (score >90 percentile) or validated target among differentially expressed protein-coding genes and the miRNA and mRNA showed inverse expression consistent with miRNA-mediated repression. We constructed broader M1 networks and temporally filtered networks, then prioritized the most stringent network consisting of temporally dynamic miRNAs and directionally reversing protein-coding genes. Networks were visualized in Cytoscape.^32^ Detailed database sources, filtering criteria, and hub definitions are provided in the **Supplementary Methods**.

High-confidence miRNA–mRNA pairs were selected based on three primary criteria: (1) differential expression of both the miRNA and target mRNA at both M1 and M2; (2) placental expression of the miRNA or target confirmed in at least one database; (3) published literature support for PE association or functional relevance to trophoblast biology. Selected pairs were additionally annotated for mechanistic connections to VGLL3/Plazyo placental findings to provide biological context for cross-compartment pathway convergence, though this annotation was not used as an exclusionary criterion. Full selection criteria, database sources, and VGLL3 connection annotations are provided in the **Supplementary Methods** and **Supplementary Table S6**.

### Placental Expression Assessment

To assess whether selected blood-derived miRNA–mRNA pairs were expressed in placental tissue, we evaluated internal VDAART placental RNA-seq data and public expression resources. Internal placental datasets included miRNA-seq from snap-frozen villous tissue and mRNA-seq from FFPE villous tissue collected from uncomplicated term pregnancies.^33^ Public resources included the Human Protein Atlas for mRNA/protein expression and miTED for miRNA expression.^34,35^ Expression evidence was compiled for temporally dynamic miRNAs and target genes in the stringent miRNA–mRNA network. Because these datasets were used to assess expression plausibility rather than disease-specific placental differential expression, placental results were interpreted as supportive biological context. Detailed placental processing and expression assessment methods are provided in the **Supplementary Methods**.

### Sensitivity Analyses

We performed sensitivity analyses to evaluate whether the main differential expression and temporal signatures were influenced by maternal vitamin D levels, PE severity, or estimated leukocyte composition. First, differential expression models were repeated with maternal 25(OH)D concentrations at early and late pregnancy included as an additional covariate. Second, severe and nonsevere PE cases were compared in exploratory analyses. Third, relative leukocyte fractions were estimated using CIBERSORT through the immunedeconv R package,^36,37^ and differential expression analyses were repeated with adjustment for cell-composition structure. Results from sensitivity models were compared with primary models using effect-size correlations and overlap of significant features. Detailed deconvolution settings, vitamin D measurement and sensitivity-analysis procedures are provided in the **Supplementary Methods.**

### Experimental Validation of miR-101-3p Regulation of DEPP1

To evaluate the predicted regulatory relationship between hsa-miR-101-3p with both *DEPP1* and *ADGRA2*, we performed transient miR-101-3p overexpression studies in HTR-8/SVneo trophoblast cells. These pairs were selected based on temporal biphasic, inverse miRNA–mRNA patterning, target-database support, placental expression evidence, and biological relevance to FOXO-related oxidative stress and autophagy programs for *DEPP1*, and angiogenesis and trophoblast-related signaling for *ADGRA2* (**Supplementary Tables: Table S8**).

HTR-8/SVneo cells were transfected with hsa-miR-101-3p mimic or negative control mimic and harvested at 24 and 48 hours after transfection. miR-101-3p expression, DEPP1 mRNA abundance, and ADGRA2 mRNA abundance were quantified using TaqMan qPCR. DEPP1 protein abundance was quantified usingWestern blotting. Relative miRNA and mRNA expression were calculated using the 2(−ΔΔCt) method, normalized to RNU48 and ACTB, respectively. DEPP1 protein abundance was normalized to α-actinin. Three independent experiments were performed. Between-group comparisons at each timepoint were evaluated using two-tailed unpaired t-tests, and two-way ANOVA was used to evaluate treatment, time, and interaction effects. Detailed cell culture, transfection, qPCR, Western blotting, antibody, and imaging procedures are provided in the **Supplementary Methods**.

### Software and Reproducibility

All statistical analyses were performed in R. Key packages included DESeq2, multiMiR, GSVA, clusterProfiler, immunedeconv, and ggplot2. Cytoscape was used for network visualization, and GraphPad Prism was used for experimental validation analyses. Unless otherwise specified, tests were two-sided, and FDR control was applied where appropriate. Detailed software versions are provided in the **Supplementary Methods**. Please see the Major Resources Table in the Supplemental Materials for a complete list of reagents, databases, software, and biological resources used in this study.

## RESULTS

### Baseline Characteristics and RNA-seq Quality Control

**Figure 1** summarizes the study analytical workflow. The study cohort comprised 43 women who developed PE and 106 normotensive controls selected from VDAART (**Supplementary Tables: Table S1**). Cases and controls were well matched on maternal age, race/ethnicity, clinical site, and treatment arm. PE cases had lower early-pregnancy 25-hydroxyvitamin D [25(OH)D] concentrations and higher body mass index (BMI) at enrollment. BMI was included as a covariate in the primary differential expression models, and 25(OH)D was evaluated in sensitivity analyses given its baseline difference between groups and correlation with BMI. Among PE cases, 11 (25.6%) had severe features, and gestational age at delivery was significantly earlier in PE cases than in controls.

Following quality control, 258 whole-blood RNA-seq samples were analyzed: 123 from early pregnancy (M1: 41 PE cases, 82 controls) and 135 from late pregnancy (M2: 38 PE cases, 97 controls), with 109 participants contributing paired samples across both timepoints (36 PE cases and 73 controls; **Supplementary Tables: Table S1**). After exclusion of sex chromosome features and low-abundance transcripts, 53,610 autosomal genes and 2,054 miRNAs were retained for analysis. Principal component analysis (PCA) showed substantial overlap between PE cases and controls. Because heme-metabolism pathways emerged in downstream analyses, we evaluated erythroid and hemoglobin signatures; neither differed significantly between PE cases and controls at either timepoint, arguing against differential red blood cell contamination as the explanation for the heme-related findings (**Supplementary Tables: Tables S2, S3, and S4**).

### Early Pregnancy Preeclampsia Signatures Show Inverse miRNA–mRNA Regulation

At early pregnancy (M1: 10–18 weeks’ gestation), differential expression analysis identified 238 mRNA-seq features associated with subsequent PE (FDR < 0.05 and |log₂FC| > 0.5), including 66 protein-coding genes, 120 long noncoding RNAs. Because the primary objective was to define miRNA-mediated regulatory programs and pathway-level mechanisms, downstream pathway and target-integration analyses focused on protein-coding transcripts and differentially expressed miRNAs.

Early pregnancy dysregulation showed marked directional asymmetry. Among the 238 differentially expressed mRNA-seq features, 229 (96.2%) were downregulated and only 9 (3.8%) were upregulated in PE cases relative to controls. This pattern was maintained among protein-coding genes, with 62 downregulated and 4 upregulated. Notable downregulated genes included ***DEPP1****, **HSD17B1**, **UHRF1**, **MMRN2**, and **SERPIND1***, implicating stress-response, placental hormone biosynthesis, epigenetic regulation, vascular remodeling, and coagulation-related pathways.

In contrast, 65 miRNAs were differentially expressed at M1, with the opposite directionality: 63 (96.9%) were upregulated and 2 (3.1%) were downregulated in PE cases. Among the most significant miRNAs were **miR-101-3p**, **miR-21-5p**, **miR-15a-5p**, **miR-16-5p**, endothelial-enriched **miR-126-3p/5p**, and immune-enriched **miR-142-5p**. Thus, early pregnancy PE was characterized by near-universal miRNA upregulation coupled with near-universal gene downregulation, consistent with broad post-transcriptional regulatory dysregulation before clinical disease onset. Complete differential expression results are provided in **Supplementary Data 1**.

### Temporal Analysis Reveals Stage-Specific and Biphasic Signatures

To determine whether early molecular signatures persisted into late pregnancy, we tracked M1-dysregulated protein-coding genes and miRNAs at M2 (32–38 weeks’ gestation). Rather than showing stable same-direction dysregulation across pregnancy, most features were either M1-specific or biphasic.

Among the 66 M1-dysregulated protein-coding genes, 20 (30%) showed biphasic pattern, with all 20 shifting from downregulation at M1 to upregulation at M2. The remaining 46 genes (70%) were M1-specific, and no protein-coding gene maintained same-direction dysregulation across both timepoints. miRNA temporal patterns showed a parallel but opposite trajectory. Among the 65 M1-dysregulated miRNAs, 10 showed biphasic pattern from upregulation at M1 to downregulation at M2, one miRNA (miR-3178) showed persistent upregulation, and 54 were M1-specific.

This pattern indicates that most early PE-associated signatures were not stable markers across pregnancy but instead either resolved or reversed direction by late gestation. At M1, miRNAs were predominantly upregulated while protein-coding genes were predominantly downregulated; at M2, a subset of these features reversed direction, suggesting stage-dependent remodeling of PE-associated regulatory networks. Among M1-dysregulated mRNA-seq features with detectable expression at M2, effect sizes were strongly negatively correlated between M1 and M2 (Spearman ρ = −0.76; Pearson r = −0.83, P < 0.0001), confirming that directional reversal was coordinated rather than random. A parallel pattern was observed for temporally overlapping miRNAs: 10 of 11 showed directional reversal from upregulation at M1 to downregulation at M2, with miR-3178 representing the sole exception showing same-direction persistence. When mRNA and miRNA effect sizes were evaluated jointly across all temporally overlapping features (N = 130), the negative correlation was maintained and strengthened (Spearman ρ = −0.82; Pearson r = −0.86, P < 0.0001), further supporting coordinated biphasic reorganization across both molecular classes (**Supplementary Data 1**). This temporal pattern was notable because the biphasic features were not randomly distributed across molecular classes. Among protein-coding genes, all 20 overlapping differentially expressed genes reversed from lower expression in PE at M1 to higher expression in PE at M2, whereas no protein-coding gene showed same-direction persistence across pregnancy. Similarly, most temporally overlapping miRNAs reversed in the opposite direction, shifting from higher expression in PE at M1 to lower expression in PE at M2. This reciprocal pattern provided the rationale for prioritizing temporally dynamic miRNA–mRNA pairs in subsequent network analyses, because the paired reversal of miRNAs and target transcripts is consistent with stage-dependent remodeling of post-transcriptional regulation.

### Pathway Analysis Identifies PI3K/AKT/mTORC1 Remodeling and Progressive Heme-Metabolism Suppression

We next performed preranked GSEA using MSigDB Hallmark gene sets separately at M1 and M2 to define pathway-level temporal patterns (**Figure 2; Supplementary Table S5**). At M1, 15 Hallmark pathways were significantly enriched, with mixed directionality. Upregulated pathways included protein secretion, androgen response, cellular stress / DNA-damage–associated transcriptional program, and **PI3K/AKT/mTORC1 signaling**, whereas downregulated pathways included cell-cycle programs, DNA repair, interferon alpha response, cholesterol homeostasis, xenobiotic metabolism, estrogen response, and **heme metabolism (Figure 2A)**. This mixed pathway profile indicates broad early molecular dysregulation involving growth, immune, proliferative, and metabolic programs.

**Figure 2.**
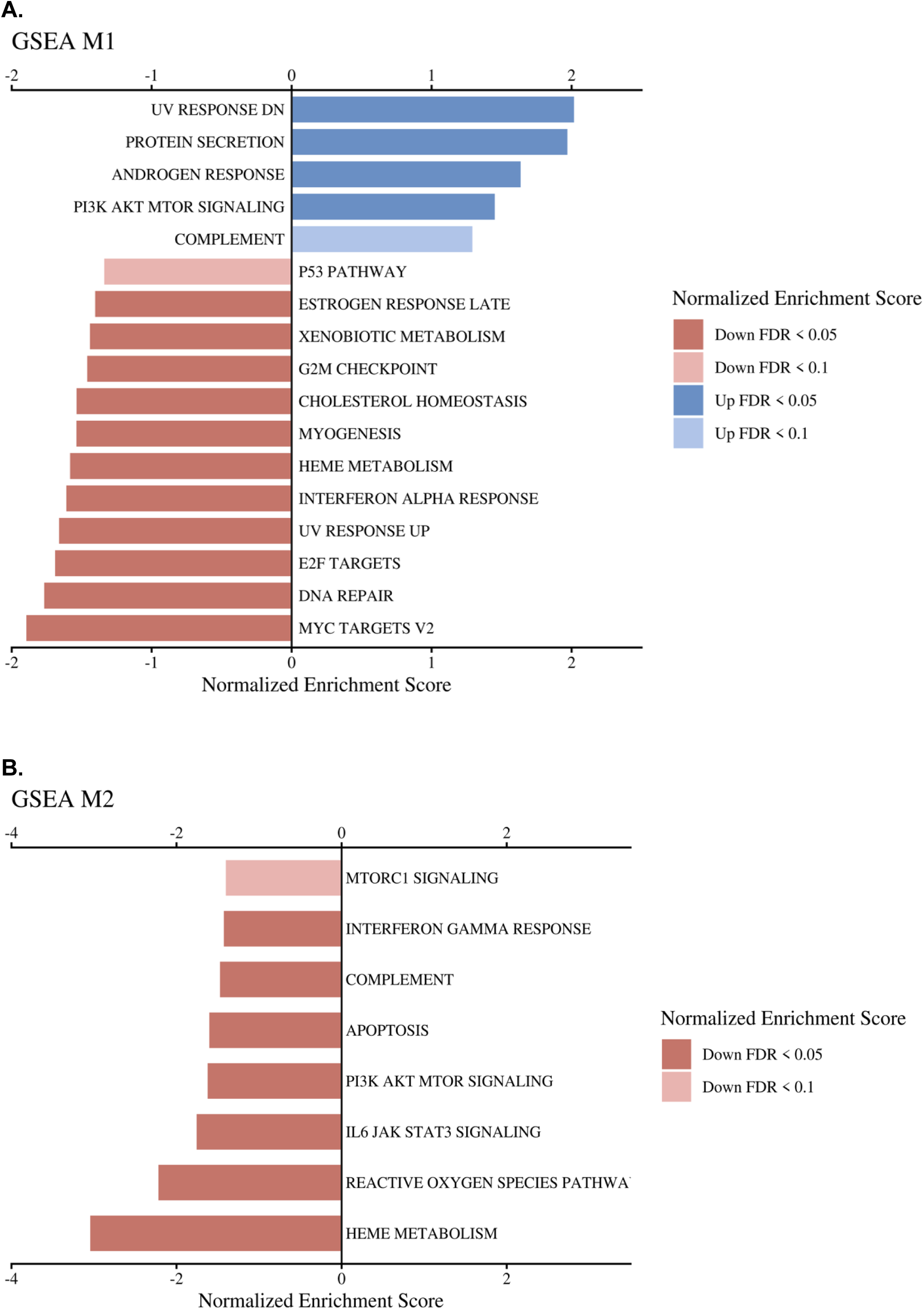
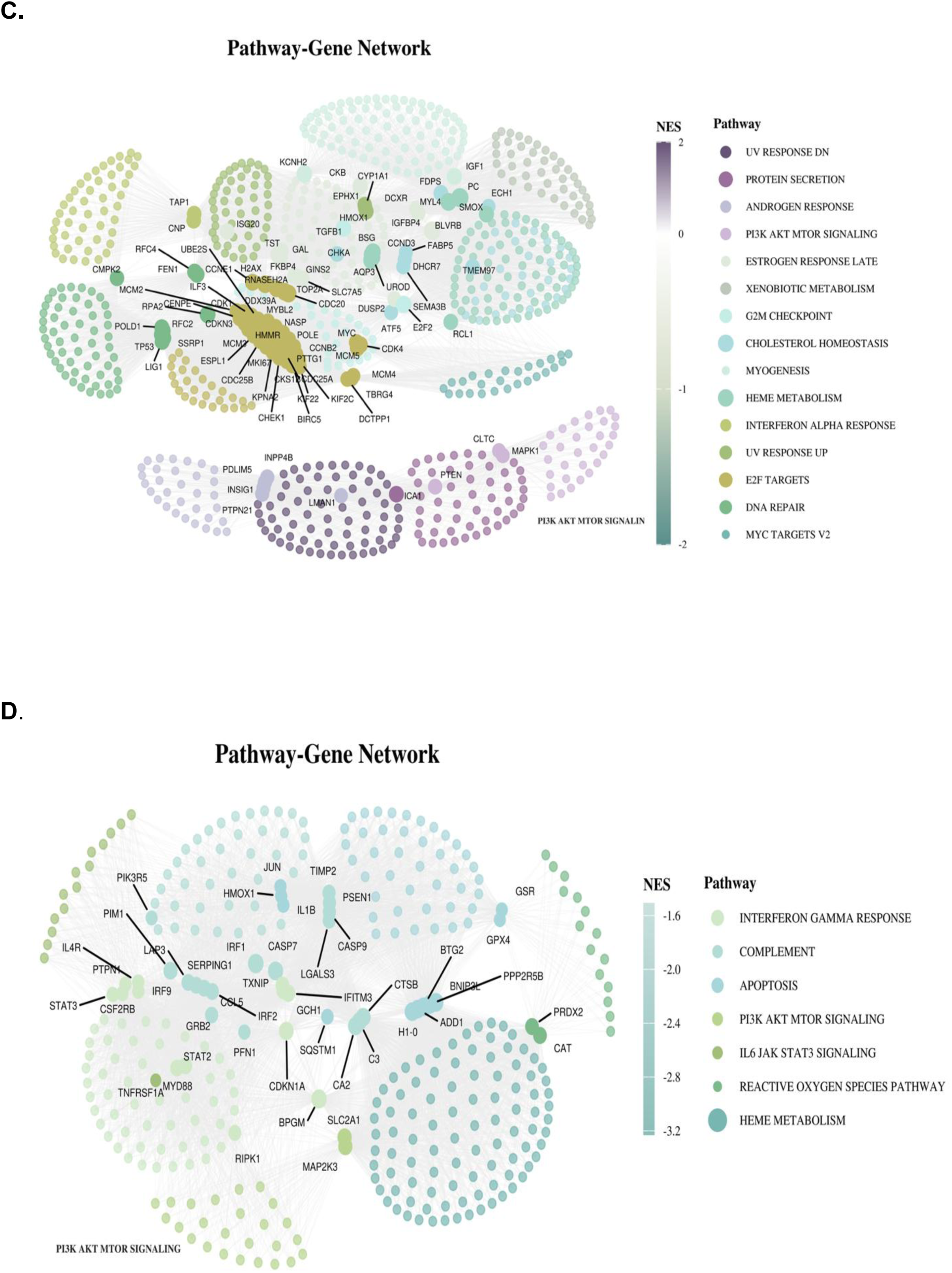
Gene Set Enrichment Analysis Reveals Temporal Pathway Dynamics in Preeclampsia (A) GSEA M1 (Early Pregnancy, 10-18 weeks). Gene Set Enrichment Analysis (GSEA) showing significantly enriched pathways in maternal whole blood from women with preeclampsia compared to controls at early pregnancy (M1). Blue bars indicate upregulated pathways (positive normalized enrichment score, NES); red/pink bars indicate downregulated pathways (negative NES). Color intensity reflects statistical significance (FDR < 0.05, darker; FDR < 0.1, lighter). At M1, preeclamptic blood exhibits activation of PI3K/AKT/mTOR signaling, protein secretion, and androgen response pathways, while showing suppression of DNA damage response, cell cycle checkpoints (G2M, E2F targets), metabolic pathways (heme metabolism, cholesterol homeostasis, xenobiotic metabolism), and stress response pathways (p53, interferon alpha, UV response). **(B) GSEA M2 (Late Pregnancy, 32-38 weeks).** Gene Set Enrichment Analysis showing significantly enriched pathways in maternal whole blood from women with preeclampsia compared to controls at late pregnancy (M2). Red/pink bars indicate downregulated pathways (negative normalized enrichment score, NES). Color intensity reflects statistical significance (FDR < 0.05, darker; FDR < 0.1, lighter). At M2, preeclamptic blood exhibits widespread pathway suppression, including dramatic downregulation of mTORC1 signaling, PI3K/AKT/mTOR signaling, immune response pathways (interferon gamma response, IL-6/JAK/STAT3 signaling, complement), apoptosis, reactive oxygen species (ROS) pathways, and heme metabolism. This pattern represents a striking reversal from M1, where PI3K/AKT/mTOR and complement pathways were upregulated, suggesting temporal remodeling of systemic signaling networks across pregnancy progression in preeclampsia. **(C) M1 Pathway-Gene Network (Early Pregnancy, 10-18 weeks).** Pathway-gene network showing differentially enriched hallmark pathways and their constituent genes in preeclamptic maternal blood at early pregnancy (M1). Each node represents a gene, with genes clustered by pathway membership. Node color indicates the normalized enrichment score (NES) of the pathway to which each gene belongs (purple/blue: upregulated pathways, NES > 0; green/teal: downregulated pathways, NES < 0). Labeled genes highlight key regulators within each pathway cluster. At M1, PI3K/AKT/mTOR signaling (purple cluster, left) and protein secretion pathways show positive enrichment, while DNA repair, cell cycle (E2F targets, MYC targets), and metabolic pathways (heme metabolism) show negative enrichment. Network layout spatially separates upregulated from downregulated pathway modules. **(D) M2 Pathway-Gene Network (Late Pregnancy, 32-38 weeks).** Pathway-gene network showing differentially enriched hallmark pathways and their constituent genes in preeclamptic maternal blood at late pregnancy (M2). Node color indicates pathway NES (all pathways downregulated: light green, NES ≈ -1.5; dark teal/blue, NES ≈ -3.0). At M2, widespread pathway suppression is evident across immune signaling (interferon gamma response, complement, IL-6/JAK/STAT3), apoptosis, PI3K/AKT/mTOR signaling, reactive oxygen species pathways, and heme metabolism. Notably, PI3K/AKT/mTOR signaling (light green cluster, left) shifts from upregulated at M1 (panel C) to downregulated at M2, demonstrating temporal pathway reversal. Network topology reveals coordinated suppression of interconnected stress response and nutrient sensing modules in late preeclamptic pregnancy.

At M2, fewer Hallmark pathways were significantly enriched, and all showed negative enrichment. The most strongly suppressed pathways were **heme metabolism** and **reactive oxygen species pathway**, followed by **IL6/JAK/STAT3 signaling**, **PI3K/AKT/mTORC1 signaling**, complement/interferon-related pathways, and programmed cell-death-related pathways (**Figure 2B**). Thus, late pregnancy PE was characterized by a more restricted pattern dominated by pathway suppression. The contrast between M1 and M2 pathway profiles suggested that PE-associated transcriptional programs change substantially across gestation. At M1, pathway enrichment was broader and bidirectional, with simultaneous activation of growth/secretory pathways and suppression of cell-cycle, immune, and metabolic programs. By M2, the pathway landscape became more restricted and uniformly negative, indicating a more restricted late-pregnancy pathway profile dominated by negative enrichment. This shift paralleled the gene-level temporal findings, in which most early pregnancy signatures were not maintained in the same direction at late pregnancy (**Figure 2C-D**).

Comparison across timepoints identified two major pathway patterns. First, **PI3K/AKT/mTORC1 signaling** showed biphasic pattern, with positive enrichment at M1 (NES = 1.46, FDR = 0.028) and negative enrichment at M2 (NES = −1.64, FDR = 0.025). Network-level analysis indicated that this reversal reflected changes in pathway gene composition rather than uniform reversal of the same pathway genes (**Supplementary Table S5**). Genes contributing to the M1-enriched pathway included upstream regulatory components such as *PTEN, GSK3B, SMAD2, MAPK1*, and *MAPK8*, whereas M2-enriched pathway components included AKT1, GRB2, TSC2, MYD88, and TRIB3. A subset of seven genes — including *MAPK1* and *YWHAB* — appeared in leading edges at both timepoints, serving as temporal bridges between stage-specific signaling modules, while stage-enriched hub genes (*PTEN* and *GSK3B* at M1; *AKT1* and *GRB2* at M2) highlighted distinct regulatory architectures at each stage (**Figure 3A**). Integration of temporally dynamic miRNAs with PI3K/AKT/mTORC1 pathway genes further suggested that miRNA-mediated regulation may contribute to stage-dependent pathway remodeling, with several reversing miRNAs predicted to target central PI3K pathway components (**Figure 3B**). Detailed Hallmark pathway results and PI3K/AKT/mTORC1 gene membership are provided in **Supplementary Appendix: Results 4** and **7** and **Supplementary Table S5**.

**Figure 3.**
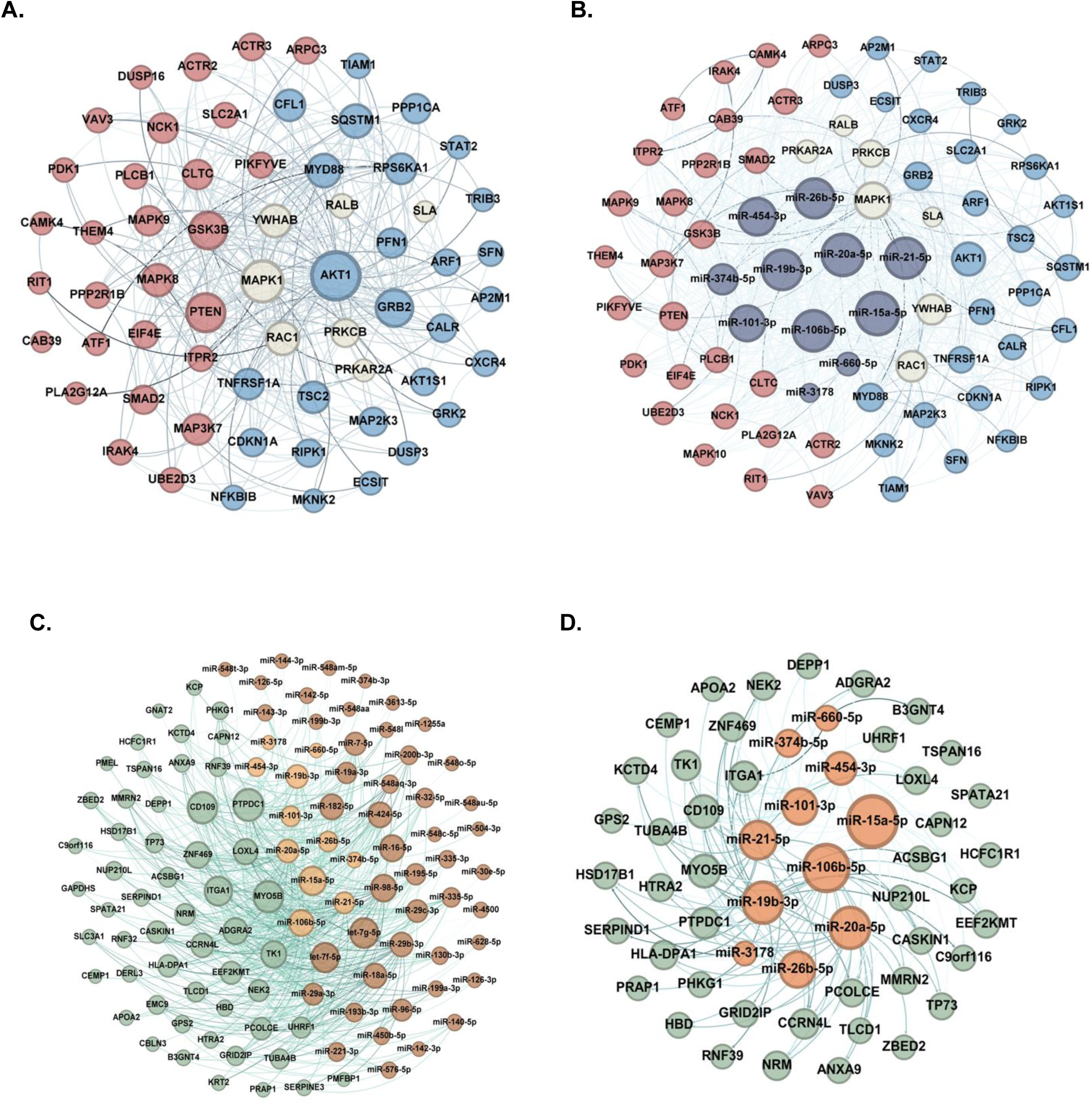
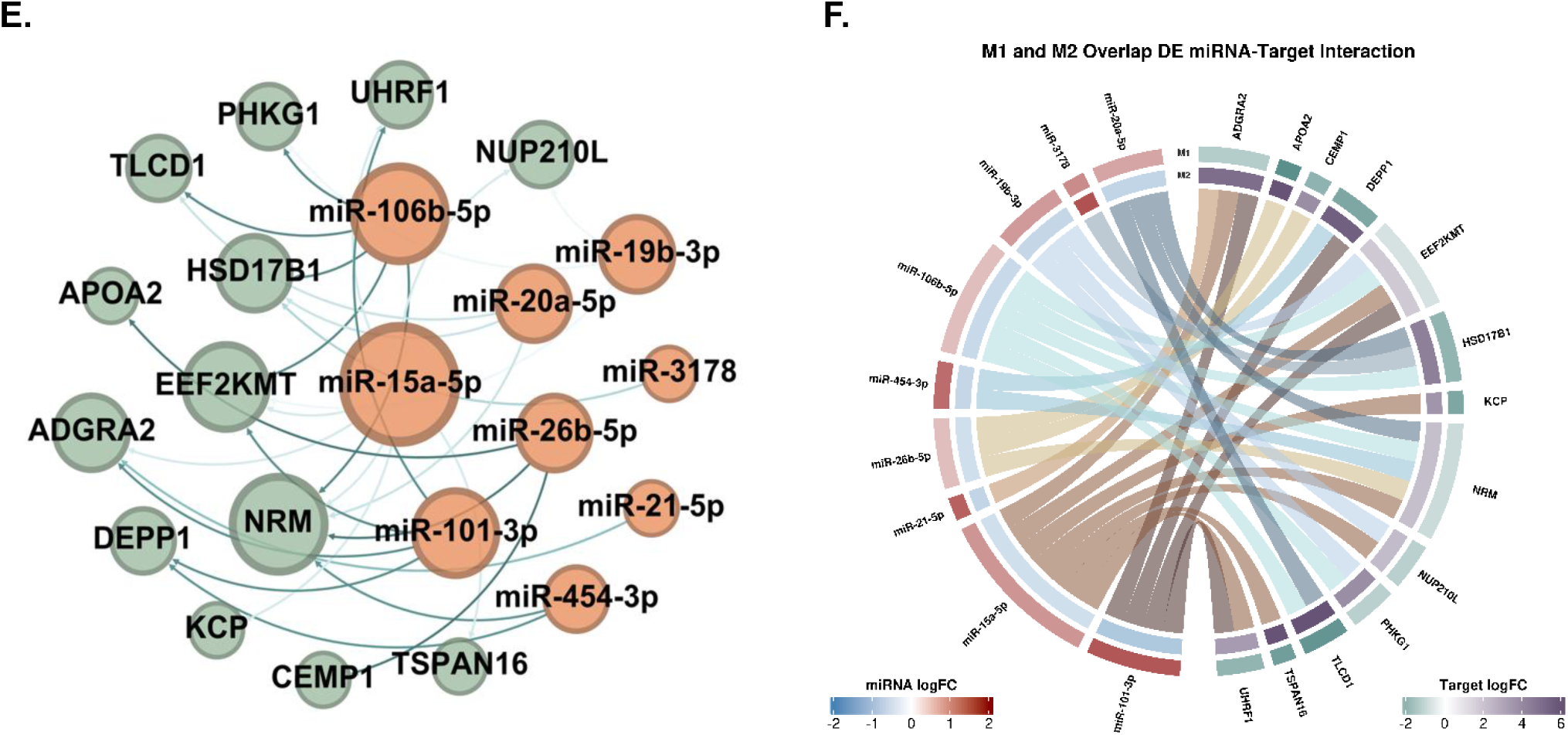
PI3K/AKT/mTORC1 Network Remodeling and miRNA Regulatory Integration Across Pregnancy in Preeclampsia A-B: PI3K/AKT/mTOR Pathway Gene Network Reveals Stage-Specific Molecular Architecture. **(A)** Gene network showing PI3K/AKT/mTOR Hallmark pathway constituents spatially organized by temporal enrichment pattern. The pathway showed directional enrichment reversal across gestation, from positive enrichment at M1 (NES = 1.46, FDR = 0.028) to negative enrichment at M2 (NES = −1.64, FDR = 0.025). Network layout reflects this temporal progression: left side (red nodes): genes enriched in the pathway leading edge at M1 only (early pregnancy, when pathway is positively enriched); center (khaki nodes, N=7): genes enriched in the pathway leading edge at both M1 and M2 (MAPK1, PRKCB, RAC1, RALB, YWHAB, PRKAR2A, and SLA), representing a core of persistently dysregulated pathway components; right side (blue nodes): genes enriched in the pathway leading edge at M2 only (late pregnancy, when pathway is negatively enriched). Node size reflects degree (number of connections). Spatial separation of M1-and M2-enriched genes demonstrates that pathway reversal involves extensive remodeling of gene composition, with distinct molecular players dominating at each pregnancy stage. MAPK1 (ERK2), the largest central node, serves as a temporal bridge connecting stage-specific pathway components. Stage-enriched hub genes further illustrate the remodeling: PTEN and GSK3B are prominent M1-specific nodes (red), while AKT1 and GRB2 are prominent M2-specific nodes (blue). **(B)** Integration of differentially expressed miRNAs onto the PI3K/AKT/mTOR pathway gene network. The protein-protein interaction network from panel A is overlaid with differentially expressed miRNAs (dark nodes) connected to their predicted target genes within the pathway. Gene nodes retain the same color scheme as panel A: red (M1-enriched genes), blue (M2-enriched genes), and khaki (N=7 genes enriched at both timepoints). miRNA nodes are positioned proximal to their predicted target genes, with edges indicating regulatory relationships. This overlay reveals that post-transcriptional regulation by miRNAs contributes to the temporal remodeling of PI3K/AKT/mTOR signaling, with stage-specific miRNAs targeting distinct subsets of pathway genes across pregnancy. Convergent targeting of central hub genes by multiple miRNAs suggests combinatorial miRNA regulation of key pathway nodes. **C-D: Temporally dynamic miRNA–mRNA regulatory networks in preeclampsia**. **(C) M1 miRNA–mRNA regulatory network.** Network showing predicted regulatory interactions between differentially expressed miRNAs and their target mRNAs in maternal blood at early pregnancy, M1, 10–18 weeks. Green nodes represent M1 differentially expressed mRNAs; orange nodes represent M1 differentially expressed miRNAs that were also differentially expressed at M2, temporally overlapping miRNAs, N=11; smaller tan nodes represent M1-specific differentially expressed miRNAs. Edges connect miRNAs to predicted target genes. Key hub genes include **DEPP1**, **ADGRA2**, **NRM**, and **CD109**; hub miRNAs include **miR-101-3p**, **miR-15a-5p**, **miR-21-5p**, and **miR-106b-5p**. Orange-highlighted miRNAs represent candidates for temporally dynamic regulatory roles because they were dysregulated at both early and late pregnancy. **(D) M1 mRNAs targeted by temporally overlapping miRNAs.** Focused regulatory network showing M1 differentially expressed mRNAs, green nodes, predicted to be targeted by the N=11 temporally overlapping differentially expressed miRNAs, orange nodes from Panel C. This subset represents early pregnancy gene-expression changes linked to miRNAs that remain dysregulated at both pregnancy stages. Central hub targets include **NRM**, **EEF2KMT**, **MYO5B**, **PTPDC1**, and **CD109**, which are regulated by multiple temporally overlapping miRNAs, including **miR-15a-5p**, **miR-20a-5p**, **miR-21-5p**, **miR-101-3p**, and **miR-106b-5p**. These interactions identify candidate regulatory relationships that may contribute to stage-dependent transcriptional remodeling across pregnancy. **(E) Stringent temporally dynamic miRNA–mRNA network with directional reversal.** The stringent network comprised 29 high-confidence miRNA–mRNA pairs involving 9 miRNAs and 13 target genes. Network depicting temporally overlapping differentially expressed miRNAs, orange nodes, targeting protein-coding mRNAs that showed directional reversal across pregnancy, green nodes. This core network represents regulatory interactions detected across both pregnancy stages, with predominantly directional reversal over time. Key prioritized regulatory pairs include **miR-101-3p → DEPP1**, **miR-101-3p → ADGRA2**, and **miR-20a-5p/miR-106b-5p → HSD17B1**. Other multi-targeting miRNAs include **miR-15a-5p**, **miR-21-5p**, **miR-26b-5p**, **miR-106b-5p**, and **miR-454-3p**. Hub mRNAs including **NRM**, **EEF2KMT**, **ADGRA2**, **HSD17B1**, and **PHKG1** receive input from multiple miRNAs, suggesting combinatorial post-transcriptional regulation. Bold edges indicate prioritized or literature-supported pairs, including **miR-101-3p → ADGRA2**, Wnt7 signaling; **miR-101-3p → DEPP1**, selected for experimental validation; and **miR-20a-5p/miR-106b-5p → HSD17B1**, estrogen biosynthesis. All pairs except **miR-3178 → HSD17B1** showed canonical directional reversal, with miRNA upregulation and target downregulation at M1 followed by miRNA downregulation and target upregulation at M2. **(F) Directional dynamics of temporally overlapping regulatory interactions.** Chord diagram illustrating the direction and magnitude of expression changes for the miRNA–mRNA regulatory pairs shown in Panel C. Outer segments represent individual miRNAs and target mRNAs, with segment size proportional to the number of interactions. Ribbons connect miRNAs to predicted target genes. Inner and outer rings show M1 and M2 expression changes, respectively. Most regulatory pairs showed canonical directional reversal: miRNAs were upregulated and their target mRNAs were downregulated at M1, followed by miRNA downregulation and target mRNA upregulation at M2. This pattern is consistent with stage-dependent remodeling of post-transcriptional regulation across pregnancy. The only non-canonical exception was **miR-3178 → HSD17B1**, in which both **miR-3178** and **HSD17B1** were upregulated at M2, consistent with regulation beyond classical miRNA-mediated mRNA degradation or with indirect regulatory effects.

Second, **heme metabolism pathway** showed progressive suppression across gestation, with negative enrichment at M1 that became substantially stronger at M2 (M1 NES = −1.56; M2 NES = −3.23), representing an approximately 2.1-fold intensification (**Figure 2A–B**). Notably, the leading-edge gene sets differed compositionally between timepoints: the M1 signature was enriched for canonical erythroid biosynthesis and differentiation genes (*HBB*, *PPOX*, *EPOR*, *TSPO2*), whereas the M2 signature additionally included heme-sensing and iron-recycling regulators (*EIF2AK1*, *NCOA4*, *SNCA*, *ISCA1*), suggesting a shift toward broader heme/iron regulatory suppression at late gestation (**Supplementary Table S5**). Importantly, erythroid and hemoglobin signature analyses did not support differential erythroid contamination or hemolysis as explanations for either signal (**Supplementary Table S4**). Together, these findings indicate progressive suppression of heme-metabolism-related transcriptional programs in maternal whole blood that intensifies and broadens mechanistically as pregnancy advances.

Broader GSEA using MSigDB immunologic (C7) and cell-type (C8) signatures showed extensive M1-specific enrichment — including immune cell activation states, T-and B-cell differentiation programs, and erythroid, NK/NKT, and endothelial cell-type signatures — but no significant C7 or C8 enrichment at M2, supporting early-stage immune and cell-type-associated dysregulation distinct from later Hallmark pathway suppression. FOXO-related immunologic signatures provided additional support for PI3K/AKT activation at M1, with gene sets representing FOXO-suppressed targets positively enriched (NES = +2.37) and FOXO target gene sets negatively enriched (NES = −2.06), consistent with FOXO transcriptional inactivation. Separately, DEPP1 appeared in multiple vascular endothelial C8 signatures that were downregulated at M1, supporting its relevance to endothelial regulatory programs in addition to its role as a FOXO-responsive gene. Full C7/C8 enrichment results, FOXO signature analyses, and PI3K-related supporting signatures are provided in **Supplementary Appendix: Results 5 and 6** and **Supplementary Data 2**.

### Temporal miRNA–mRNA Integration Prioritizes *ADGRA2*, *HSD17B1*, and *DEPP1* Regulatory Axes

To identify candidate post-transcriptional regulatory interactions, we integrated differentially expressed miRNAs with high-confidence predicted and validated target mRNAs. We constructed a series of increasingly stringent networks to distinguish broad early pregnancy regulatory architecture from temporally dynamic interactions across pregnancy.

First, we integrated all M1-dysregulated miRNAs with M1-dysregulated protein-coding genes showing inverse expression patterns consistent with miRNA-mediated repression (**Figure 3C**). This broader early pregnancy network identified multiple candidate hub miRNAs and target genes, including **miR-101-3p**, **miR-15a-5p**, **miR-21-5p**, **miR-106b-5p**, ***DEPP1****, **ADGRA2**, **NRM***, and **CD109**. We then focused on M1-dysregulated mRNAs targeted by the 11 miRNAs that remained differentially expressed at M2, generating a temporally filtered network of candidate interactions involving 9 miRNAs with evidence of dysregulation at both pregnancy stages and target genes at M1 (**Figure 3D**). At the third step, the most stringent analysis integrated the 11 temporally dynamic miRNAs with the 18 protein-coding genes that showed biphasicity across pregnancy. This yielded 29 high-confidence miRNA–mRNA pairs involving 9 miRNAs and 13 target genes (**Figure 3E–F**; **Supplementary Table S6**). Of these, 28 of 29 pairs followed the canonical inverse pattern expected for miRNA-mediated repression: miRNA upregulation with target mRNA downregulation at M1, followed by miRNA downregulation with target mRNA upregulation at M2. The sole exception was **miR-3178 → *HSD17B1***, in which both the miRNA and target were upregulated at M2, consistent with regulation beyond classical miRNA-mediated mRNA degradation or with indirect regulatory effects. From the 29 high-confidence pairs, 11 were selected for **Table 2** based on strength of biological evidence, placental expression support, and literature relevance; the remaining 18 are detailed in **Supplementary Table S6.**

**Table 1.** Post-QC sample distribution.

| **Sample Distribution** | **PE Cases** | **Normotensive Controls** | **Total** |
|---|---|---|---|
| **By Timepoint** |  |  |  |
| Early pregnancy (M1) | 41 | 82 | 123 |
| Late pregnancy (M2) | 38 | 97 | 135 |
| **By Longitudinal Pattern** |  |  |  |
| Early only (10-18 weeks) | 5 (11.6%) | 9 (8.5%) | 14 |
| Late only (32-38 weeks) | 2 (4.7%) | 24 (22.6%) | 26 |
| Both timepoints (paired) | 36 (83.7%) | 73 (68.9%) | 109 |
| **Total Unique Participants** | **43** | **106** | **149** |
| **Total RNA-seq Samples** | **79** | **179** | **258** |

**Table 2.**
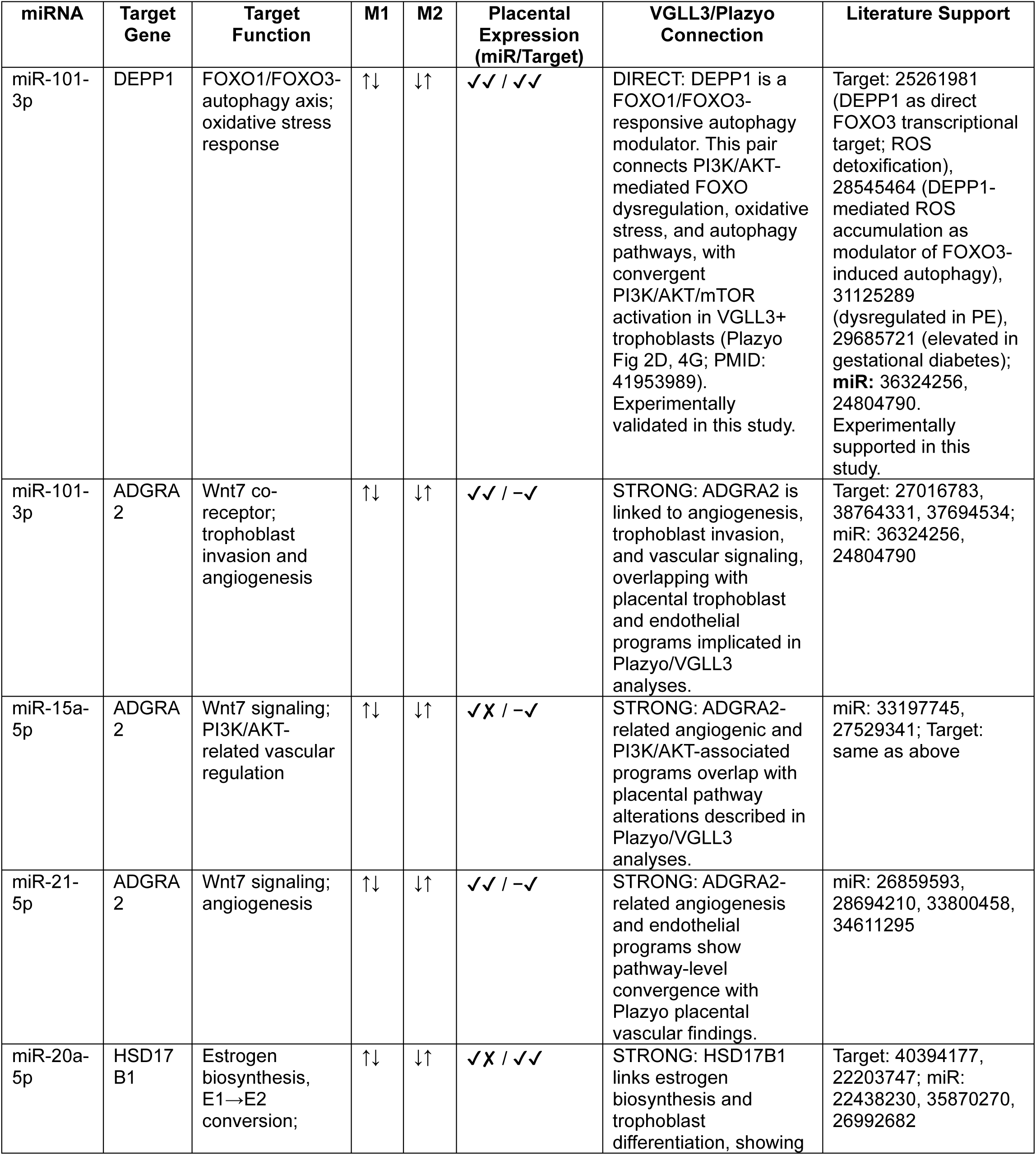
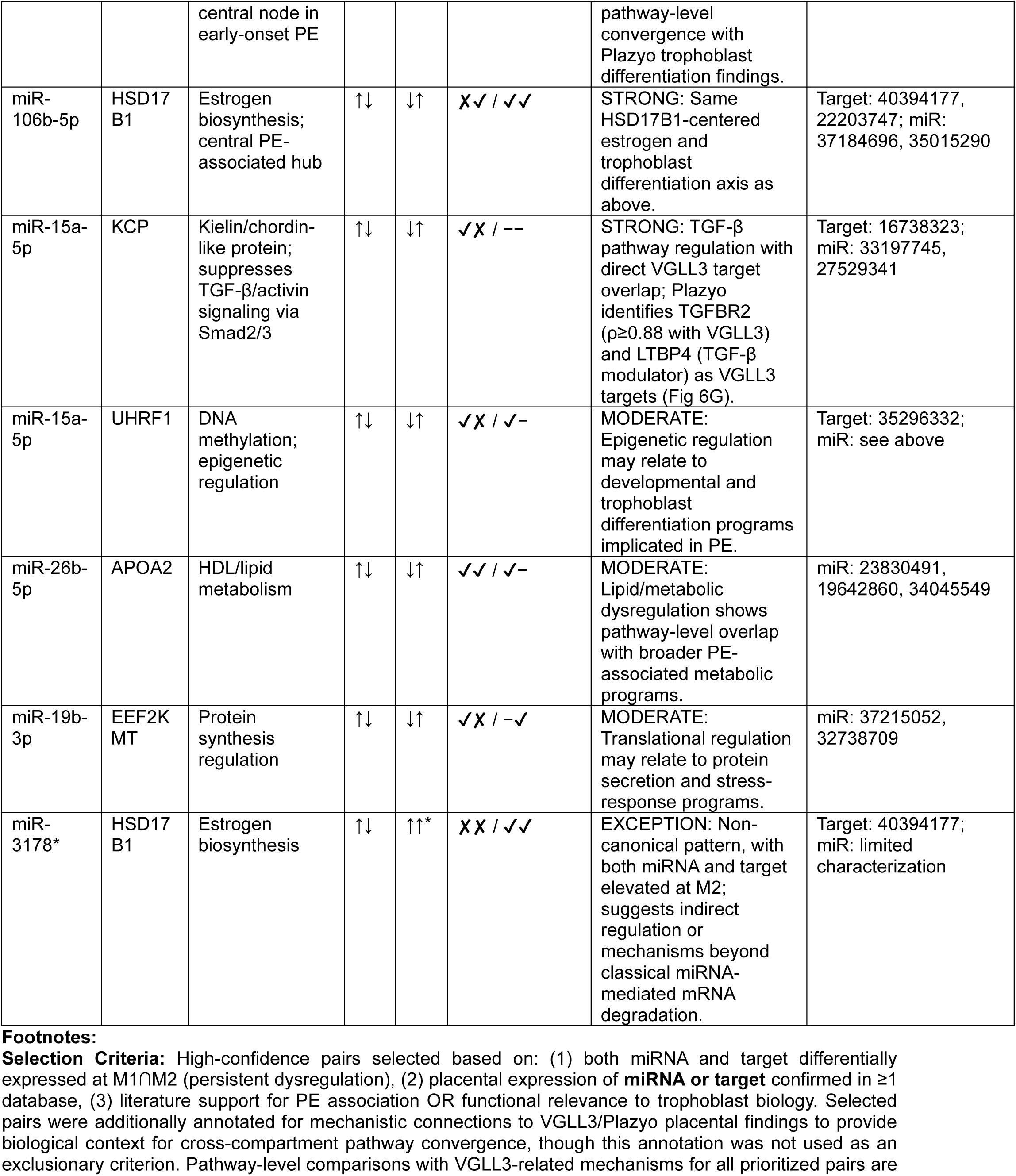
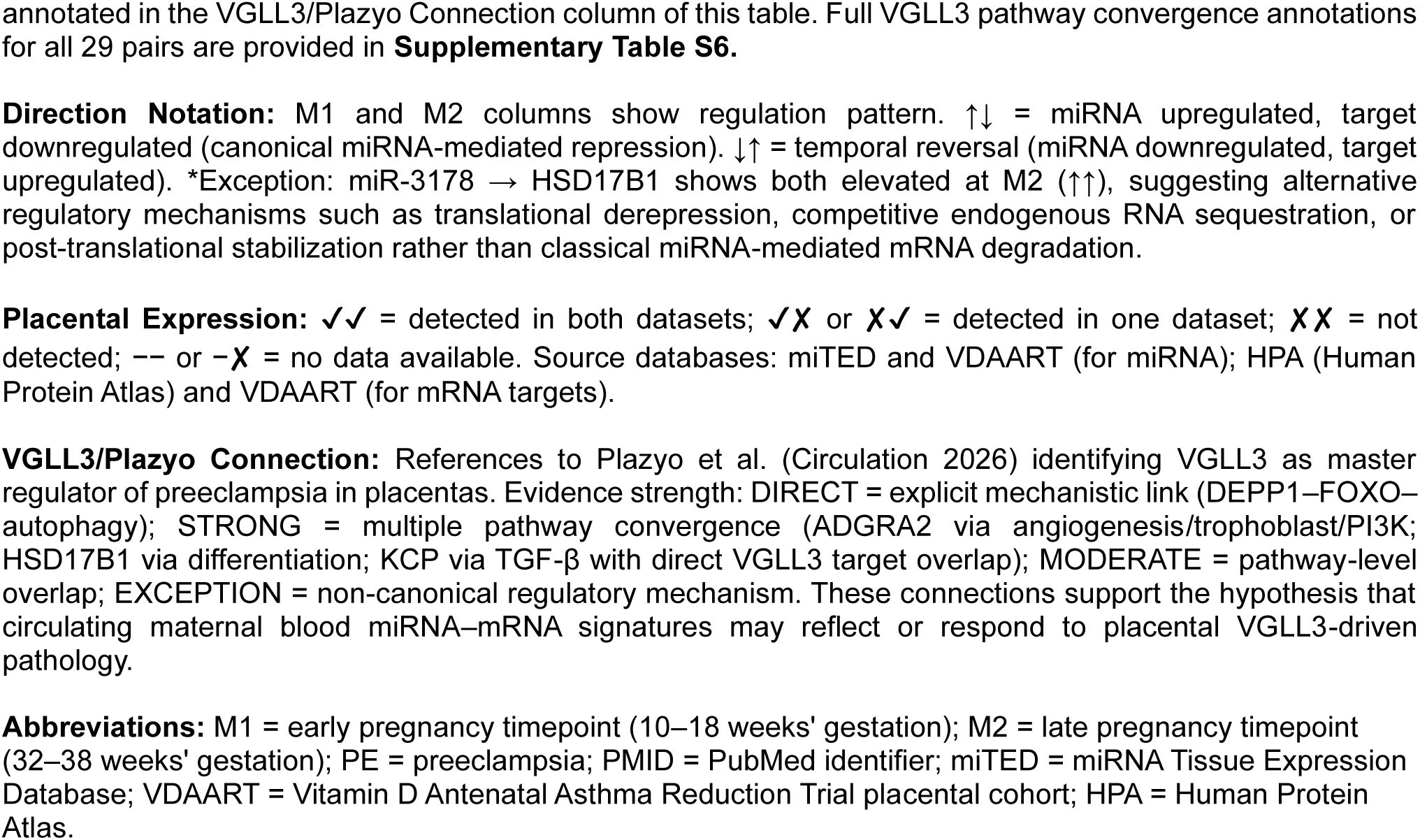
Selected high-confidence temporally dynamic miRNA–mRNA regulatory pairs in preeclampsia.

| miRNA | Target Gene | Target Function | M1 | M2 | Placental Expression (miR/Target) | VGLL3/Plazyo Connection | Literature Support |
|---|---|---|---|---|---|---|---|
| miR-101-3p | DEPP1 | FOXO1/FOXO3-autophagy axis; oxidative stress response | ↑↓ | ↓↑ | ✓✓ / ✓✓ | DIRECT: DEPP1 is a FOXO1/FOXO3-responsive autophagy modulator. This pair connects PI3K/AKT-mediated FOXO dysregulation, oxidative stress, and autophagy pathways, with convergent PI3K/AKT/mTOR activation in VGLL3+ trophoblasts (Plazyo Fig 2D, 4G; PMID: 41953989). Experimentally validated in this study. | Target: 25261981 (DEPP1 as direct FOXO3 transcriptional target; ROS detoxification), 28545464 (DEPP1-mediated ROS accumulation as modulator of FOXO3-induced autophagy), 31125289 (dysregulated in PE), 29685721 (elevated in gestational diabetes); **miR:** 36324256, 24804790. Experimentally supported in this study. |
| miR-101-3p | ADGRA2 | Wnt7 co-receptor; trophoblast invasion and angiogenesis | ↑↓ | ↓↑ | ✓✓ / −✓ | STRONG: ADGRA2 is linked to angiogenesis, trophoblast invasion, and vascular signaling, overlapping with placental trophoblast and endothelial programs implicated in Plazyo/VGLL3 analyses. | Target: 27016783, 38764331, 37694534; miR: 36324256, 24804790 |
| miR-15a-5p | ADGRA2 | Wnt7 signaling; PI3K/AKT-related vascular regulation | ↑↓ | ↓↑ | ✓✗ / −✓ | STRONG: ADGRA2-related angiogenic and PI3K/AKT-associated programs overlap with placental pathway alterations described in Plazyo/VGLL3 analyses. | miR: 33197745, 27529341; Target: same as above |
| miR-21-5p | ADGRA2 | Wnt7 signaling; angiogenesis | ↑↓ | ↓↑ | ✓✓ / −✓ | STRONG: ADGRA2-related angiogenesis and endothelial programs show pathway-level convergence with Plazyo placental vascular findings. | miR: 26859593, 28694210, 33800458, 34611295 |
| miR-20a-5p | HSD17B1 | Estrogen biosynthesis, E1→E2 conversion; central node in early-onset PE | ↑↓ | ↓↑ | ✓✗ / ✓✓ | STRONG: HSD17B1 links estrogen biosynthesis and trophoblast differentiation, showing pathway-level convergence with Plazyo trophoblast differentiation findings. | Target: 40394177, 22203747; miR: 22438230, 35870270, 26992682 |
| miR-106b-5p | HSD17B1 | Estrogen biosynthesis; central PE-associated hub | ↑↓ | ↓↑ | ✗✓ / ✓✓ | STRONG: Same HSD17B1-centered estrogen and trophoblast differentiation axis as above. | Target: 40394177, 22203747; miR: 37184696, 35015290 |
| miR-15a-5p | KCP | Kielin/chordin-like protein; suppresses TGF-β/activin signaling via Smad2/3 | ↑↓ | ↓↑ | ✓✗ / −− | STRONG: TGF-β pathway regulation with direct VGLL3 target overlap; Plazyo identifies TGFBR2 (ρ≥0.88 with VGLL3) and LTBP4 (TGF-β modulator) as VGLL3 targets (Fig 6G). | Target: 16738323; miR: 33197745, 27529341 |
| miR-15a-5p | UHRF1 | DNA methylation; epigenetic regulation | ↑↓ | ↓↑ | ✓✗ / ✓− | MODERATE: Epigenetic regulation may relate to developmental and trophoblast differentiation programs implicated in PE. | Target: 35296332; miR: see above |
| miR-26b-5p | APOA2 | HDL/lipid metabolism | ↑↓ | ↓↑ | ✓✓ / ✓− | MODERATE: Lipid/metabolic dysregulation shows pathway-level overlap with broader PE-associated metabolic programs. | miR: 23830491, 19642860, 34045549 |
| miR-19b-3p | EEF2KMT | Protein synthesis regulation | ↑↓ | ↓↑ | ✓✗ / −✓ | MODERATE: Translational regulation may relate to protein secretion and stress-response programs. | miR: 37215052, 32738709 |
| miR-3178* | HSD17B1 | Estrogen biosynthesis | ↑↓ | ↑↑* | ✗✗ / ✓✓ | EXCEPTION: Non-canonical pattern, with both miRNA and target elevated at M2; suggests indirect regulation or mechanisms beyond classical miRNA-mediated mRNA degradation. | Target: 40394177; miR: limited characterization |

**Footnotes:**
**Selection Criteria:** High-confidence pairs selected based on: (1) both miRNA and target differentially expressed at M1∩M2 (persistent dysregulation), (2) placental expression of **miRNA or target** confirmed in ≥1 database, (3) literature support for PE association OR functional relevance to trophoblast biology. Selected pairs were additionally annotated for mechanistic connections to VGLL3/Plazyo placental findings to provide biological context for cross-compartment pathway convergence, though this annotation was not used as an exclusionary criterion. Pathway-level comparisons with VGLL3-related mechanisms for all prioritized pairs are annotated in the VGLL3/Plazyo Connection column of this table. Full VGLL3 pathway convergence annotations for all 29 pairs are provided in **Supplementary Table S6.**
**Direction Notation:** M1 and M2 columns show regulation pattern. ↑↓ = miRNA upregulated, target downregulated (canonical miRNA-mediated repression). ↓↑ = temporal reversal (miRNA downregulated, target upregulated). \*Exception: miR-3178 → HSD17B1 shows both elevated at M2 (↑↑), suggesting alternative regulatory mechanisms such as translational derepression, competitive endogenous RNA sequestration, or post-translational stabilization rather than classical miRNA-mediated mRNA degradation.
**Placental Expression:** ✓✓ = detected in both datasets; ✓✗ or ✗✓ = detected in one dataset; ✗✗ = not detected; −− or −✗ = no data available. Source databases: miTED and VDAART (for miRNA); HPA (Human Protein Atlas) and VDAART (for mRNA targets).
**VGLL3/Plazyo Connection:** References to Plazyo et al. (Circulation 2026) identifying VGLL3 as master regulator of preeclampsia in placentas. Evidence strength: DIRECT = explicit mechanistic link (DEPP1–FOXO–autophagy); STRONG = multiple pathway convergence (ADGRA2 via angiogenesis/trophoblast/PI3K; HSD17B1 via differentiation; KCP via TGF-β with direct VGLL3 target overlap); MODERATE = pathway-level overlap; EXCEPTION = non-canonical regulatory mechanism. These connections support the hypothesis that circulating maternal blood miRNA–mRNA signatures may reflect or respond to placental VGLL3-driven pathology.
**Abbreviations:** M1 = early pregnancy timepoint (10–18 weeks' gestation); M2 = late pregnancy timepoint (32–38 weeks' gestation); PE = preeclampsia; PMID = PubMed identifier; miTED = miRNA Tissue Expression Database; VDAART = Vitamin D Antenatal Asthma Reduction Trial placental cohort; HPA = Human Protein Atlas.

The predominance of canonical inverse reversal distinguished this temporally dynamic network from a simple overlap of differentially expressed features. At M1, the dominant pattern was miRNA upregulation with target mRNA downregulation, whereas at M2 the same regulatory pairs showed the opposite configuration. This structure supports the interpretation that the network captures coordinated regulatory transitions across pregnancy rather than independent cross-sectional signatures at each timepoint.

The 29-pair network also showed evidence of combinatorial regulation. **miR-15a-5p** was the most connected miRNA, targeting 7 genes, while **miR-106b-5p**, **miR-101-3p**, and **miR-20a-5p** also targeted multiple temporally dynamic transcripts. Several target genes were co-regulated by multiple miRNAs, including ***NRM****, **EEF2KMT**, **ADGRA2**, **HSD17B1**, and **PHKG1***. The strongest prioritized axes included convergent targeting of ***ADGRA2*** by **miR-101-3p**, **miR-15a-5p**, and **miR-21-5p**; targeting of **H*SD17B1*** by **miR-20a-5p**, **miR-106b-5p**, and **miR-3178**; and the **miR-101-3p → *DEPP1*** pair linking temporal miRNA–mRNA reversal to the PI3K/AKT–FOXO stress-response axis. Additional prioritized pairs included **miR-15a-5p → *KCP***, **miR-15a-5p → *UHRF1***, and **miR-26b-5p → *APOA2***. Detailed literature support, placental-expression evidence, and mechanistic annotations for prioritized and full miRNA–mRNA pairs are provided in **Table 2** and **Supplementary Table S6**.

### Placental Expression Supports Biological Plausibility of Selected Blood-Derived Pairs

To assess whether selected blood-derived miRNA–mRNA pairs were biologically plausible in placental tissue, we evaluated internal VDAART placental RNA-seq data and public expression resources.^33,34^ Most temporally dynamic miRNAs showed detectable placental expression in at least one placental expression resource, although expression levels varied across datasets. Among the 13 target genes in the high-confidence 29-pair network, 8 were detected in VDAART placental RNA-seq and 5 showed detectable placental expression in the Human Protein Atlas (HPA); in total, 11 of 13 target genes were detected in at least one placental expression resource. Overall, 26 of 29 miRNA–mRNA pairs showed detectable expression of both partners in at least one placental expression resource. The strongest support was observed for **miR-101-3p → *DEPP1***, ***ADGRA2***-targeting pairs, and ***HSD17B1***-targeting pairs. These pairs were notable because they linked the blood-derived temporal network to biologically relevant placental programs: ***DEPP1*** to FOXO-mediated stress and autophagy responses, ***ADGRA2*** to angiogenic and trophoblast-related signaling, and ***HSD17B1*** to placental steroid biosynthesis and trophoblast differentiation. In contrast, some pairs showed weaker placental support, suggesting that a subset of the maternal blood network may reflect systemic maternal responses rather than placental expression programs. These findings support the biological plausibility that a subset of blood-derived regulatory signatures may reflect placental or shared maternal–placental regulatory processes, while other pairs may represent maternal systemic responses to PE. Detailed placental expression results are provided in **Supplementary Appendix Figure S1** and **Supplementary Tables: Table S6**.

### Sensitivity Analyses Support Robustness of Temporal Signatures

Sensitivity analyses supported the robustness of the temporal signatures. Temporally dynamic protein-coding genes and miRNAs did not differ significantly between severe and nonsevere PE cases, although severity-stratified analyses were limited by sample size. Adjustment for estimated leukocyte composition produced highly concordant effect sizes compared with the primary models, and no major leukocyte population differed significantly between PE cases and controls after correction for multiple testing. Adjustment for maternal 25(OH)D concentrations also yielded highly concordant differential expression estimates. Together, these analyses indicate that the main temporal signatures were not explained by estimated leukocyte composition, vitamin D status, or PE severity. Full sensitivity analyses are provided in **Supplementary Appendix: Results 11** and **Supplementary Tables: Tables S7**.

### miR-101-3p Overexpression Suppresses DEPP1 in Trophoblast Cells

We selected **miR-101-3p → *DEPP1*** and ***ADGRA2*** for functional validation because they showed coordinated temporal reversal, target-database support, placental expression, and endothelial localization. In addition, DEPP1 is connected to the PI3K/AKT–FOXO axis and ***ADGRA2*** to angiogenesis and trophoblast-related signaling (**Supplementary Tables: Table S6**). To test whether miR-101-3p can suppress DEPP1 or ADGRA2 expression in trophoblast cells, we transiently overexpressed hsa-miR-101-3p in HTR-8/SVneo cells and quantified miR-101-3p, DEPP1, and ADGRA2 mRNA at 24 and 48 hours post-transfection.

Transfection with hsa-miR-101-3p mimic resulted in marked miR-101-3p overexpression at both timepoints compared with negative control-transfected cells (24h: P = 0.008; 48h: P = 0.005; **Figure 4A**). Relative miR-101-3p expression increased approximately 800-fold at 24 hours and 300-fold at 48 hours. miR-101-3p overexpression significantly reduced **DEPP1** transcript abundance at both 24 hours (43% reduction, P < 0.001) and 48 hours (39% reduction, P = 0.007; **Figure 4B**). A smaller reduction in ADGRA2 transcript abundance was also observed at both 24 hours (18% reduction, P = 0.03) and 48 hours (29% reduction, P = 0.008; **Supplementary Appendix Figure S2**).

**Figure 4.**
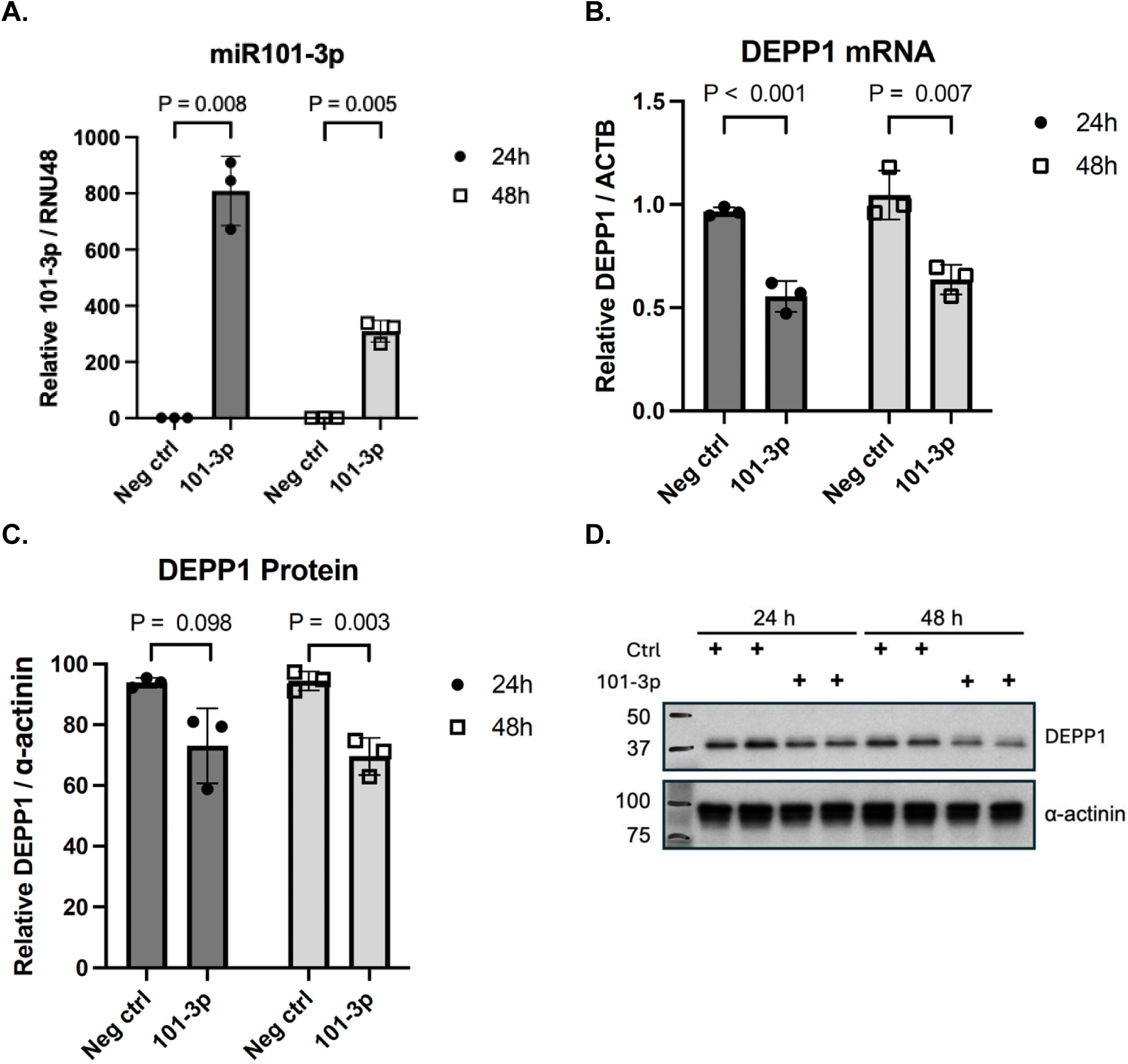
miR-101-3p overexpression suppresses DEPP1 mRNA and protein expression in HTR-8/SVneo trophoblast cells. HTR-8/SVneo cells were transfected with hsa-miR-101-3p mimic or negative control mimic and harvested at 24 or 48 hours post-transfection. (A) Relative miR-101-3p expression normalized to RNU48. (B) Relative DEPP1 mRNA expression, normalized to ACTB, was reduced at both 24 hours (43% reduction, P < 0.001) and 48 hours (39% reduction, P = 0.007) compared with the negative control. (C) DEPP1 protein abundance, normalized to α-actinin and quantified by densitometry, was reduced at 24 hours (22% reduction, P = 0.098) and 48 hours (26% reduction, P = 0.003). (D) Representative Western blot. Ctrl, negative control; 101-3p, miR-101-3p mimic. Each condition shows two biological replicates per timepoint. Data are mean ± SEM (n=3 independent experiments). Statistical comparisons used two-tailed unpaired t-tests. P values are shown.

Given the more pronounced effect of miR-101-3p overexpression on DEPP1 mRNA, we further quantified DEPP1 protein expression at 24 and 48 hours post-transfection. Consistent with the mRNA findings, DEPP1 protein abundance was also reduced following miR-101-3p overexpression (**Figure 4C–D**), with protein levels decreasing by 22% at 24 hours (P = 0.098) and by 26% at 48 hours; the reduction reached statistical significance at 48 hours (P = 0.003). The delayed protein response relative to mRNA suppression is consistent with expected protein turnover following miRNA-mediated transcript repression. Together, these results support the capacity of miR-101-3p overexpression to suppress DEPP1 mRNA and protein abundance in trophoblast cells, consistent with the predicted regulatory relationship identified from the longitudinal whole-blood miRNA–mRNA network.

## DISCUSSION

In this longitudinal RNA-sequencing study of maternal whole blood across pregnancy, we found that PE was associated with marked temporal reorganization of miRNA–mRNA regulatory networks. This pattern — characterized by early pregnancy upregulation giving way to late pregnancy downregulation of the same regulatory pairs — is consistent with the observation that PE-associated molecular signatures in maternal blood are strongly influenced by gestational stage and may depart from the predominantly linear transcriptome progressions observed in normal gestation.^38–41^

Under our prespecified differential-expression criteria, we observed minimal evidence of same-direction temporal conservation, with no protein-coding genes and only one miRNA, miR-3178, maintaining the same direction of dysregulation across both timepoints. Instead, PE-associated signatures were predominantly stage-specific or biphasic. Early pregnancy was characterized by near-universal miRNA upregulation accompanied by gene downregulation, whereas a subset of these signals reversed direction in late pregnancy. At the pathway level, PI3K/AKT/mTORC1 signaling showed dynamic remodeling, while heme-metabolism-related transcriptional programs showed progressive suppression across pregnancy. Together, these findings support a model in which the molecular architecture of PE changes substantially across gestation rather than remaining fixed.

A central implication of these findings is that PE may be better understood as a temporally evolving disorder than as a condition defined by stable transcriptomic signatures. The strong negative correlation of effect sizes between early and late pregnancy supports coordinated temporal reorganization rather than random fluctuation. This may help explain inconsistencies across prior PE transcriptomic studies performed at different gestational ages,^8,42–44^ since signatures identified in early pregnancy may differ substantially from those detected in late pregnancy or at delivery. This observation has direct implications for biomarker discovery: molecular signatures for PE may need to be interpreted and validated in a gestational-age-specific framework rather than as static disease markers.

The contrasting temporal patterns observed in this study suggest that different biological programs may operate on different timescales during PE progression. PI3K/AKT/mTORC1 signaling shifted from positive enrichment in early pregnancy to negative enrichment in late pregnancy, consistent with stage-dependent remodeling of growth and survival signaling. In contrast, heme-metabolism-related transcriptional programs were suppressed at both timepoints, with stronger suppression in late pregnancy. While the present observational data cannot establish causality or distinguish primary drivers from secondary responses, these patterns provide a useful framework for separating dynamically remodeled regulatory programs from progressively altered metabolic transcriptional programs.

### PI3K/AKT/mTORC1 Remodeling and the FOXO–DEPP1 Axis

Among the pathways identified, PI3K/AKT/mTORC1 signaling showed the clearest evidence of temporal remodeling. This pathway is known to regulate trophoblast proliferation, invasion, migration, and cellular stress responses, and its dysregulation has been implicated in PE pathogenesis.^45–49^ Specifically, suppression of PI3K/AKT/mTORC1 signaling has been documented in preeclamptic placentas,^47^ while the diagnostic significance of reduced PI3K and mTORC1 activity in maternal serum has been demonstrated,^50^ and experimental interventions that activate this pathway ameliorate trophoblast dysfunction.^48,49,51^ This temporal pattern was supported by several orthogonal analyses. In our data, PI3K/AKT/mTORC1 signaling was positively enriched at M1 and negatively enriched at M2. Network analysis further suggested that this pattern did not reflect uniform reversal of the same genes but rather remodeling of pathway composition across pregnancy, with different subsets of pathway genes contributing at each stage.

This temporal pattern was supported by FOXO-related signatures and by the behavior of DEPP1, expressed in placental tissue in published datasets and confirmed in internal placental RNA-seq.^52^ PI3K/AKT activation inhibits FOXO transcription factors through phosphorylation-mediated cytoplasmic sequestration.^53–55^ DEPP1 is a FOXO-responsive gene involved in oxidative stress and autophagy-related programs.^52,56^ In this study, DEPP1 was downregulated at M1 and upregulated at M2, paralleling the directional shift in PI3K/AKT/mTORC1 signaling. This pattern is consistent with a model in which early PI3K/AKT activation is accompanied by suppression of FOXO target programs, including DEPP1, whereas late PI3K/AKT suppression may permit re-emergence of FOXO-responsive transcriptional activity. Because DEPP1 is a direct transcriptional target of FOXO3 and functions as a reactive oxygen species (ROS)-dependent amplifier of FOXO3-induced autophagy,^56^ its suppression during the period of peak PI3K/AKT activity is mechanistically consistent with a regulatory program in which early trophoblast proliferation and invasion are prioritized over autophagy induction. The functional significance of DEPP1 suppression is further supported by its role in regulating catalase activity: DEPP1 localizes to peroxisomes and impairs catalase-mediated hydrogen peroxide detoxification in human cell models, such that reduced DEPP1 expression — as observed at M1 in this study — would be expected to transiently preserve catalase activity and attenuate ROS accumulation.^57^ Consistent with this inference, reduced erythrocyte catalase activity has been documented in women who subsequently develop PE, measured prospectively in early pregnancy and associated with increased odds of disease, ^58^ suggesting that the early suppression of DEPP1 and its downstream effects on catalase-mediated ROS handling may reflect a transient compensatory state that becomes insufficient as pregnancy advances.

The miR-101-3p → DEPP1 regulatory axis provides a functional link between the temporal miRNA–mRNA network and this pathway-level finding. miR-101-3p was strongly upregulated at M1 and downregulated at M2, while DEPP1 showed the opposite pattern. Experimental overexpression of miR-101-3p in HTR-8/SVneo trophoblast cells reduced DEPP1 mRNA and protein abundance, supporting the capacity of miR-101-3p to repress DEPP1 in a trophoblast cell model. These data do not prove that miR-101-3p directly binds DEPP1 in vivo or that this interaction drives PE progression, but they provide functional support for one prioritized regulatory interaction emerging from the longitudinal blood network.

This finding also helps contextualize prior reports on miR-101 in PE. Previous studies have implicated miR-101 in trophoblast biology, but reported directions of dysregulation and functional consequences have varied across models and contexts.^59–61^ Our data showing miR-101-3p upregulation at M1 (log₂FC = 1.94, FDR = 1.3×10⁻⁶) aligns with the Zhao et al. findings,^59^ and our experimental validation establishes DEPP1 as a functionally relevant target in the HTR-8/SVneo trophoblast model.

Our results suggest that temporal context may be important: miR-101-3p was upregulated in early pregnancy but decreased by late pregnancy among temporally dynamic miRNAs. This emphasizes the importance of considering gestational timing when interpreting miRNA function in PE. These findings can also be contextualized against our prior cross-sectional report linking miR-182-5p upregulation to suppression of IGF1R, a high-connectivity hub gene, in maternal circulation and validated experimentally in HTR-8/SVneo trophoblast cells. ^20^ In the present longitudinal dataset, miR-182-5p was upregulated at M1 in the same direction as previously reported, but was not differentially expressed at M2. IGF1R itself did not reach significance as a differentially expressed gene at either timepoint in this RNA-seq dataset; however, several IGF-axis genes were enriched in pathway-level analyses, including *IGFBP4*, *IGF1*, *IGFBP7*, and *IGFBP2* at M1 and *IGF2R* at M2. Because IGF1R is a canonical upstream activator of PI3K/AKT signaling, these findings suggest that the IGF–PI3K/AKT axis identified previously and the PI3K/AKT–FOXO–DEPP1 axis identified here may represent overlapping or convergent components of a broader growth-factor signaling network dysregulated in PE, acting through different molecular entry points across early and late gestation. The absence of *IGF1R* itself as a significant individual feature in the current analysis may reflect platform-related differences in detection sensitivity, since the prior report used microarray-based profiling whereas the present study used RNA-sequencing on the same PAXgene whole-blood samples from an overlapping subject set. These platform-level differences in dynamic range, full-transcript versus probe-based quantification, and sensitivity to lower-abundance transcripts may explain discordant feature-level detection between the two studies, rather than indicating a true absence of IGF-related dysregulation.

### Prioritized Regulatory Axes and VGLL3–Hippo Pathway Convergence

The temporally dynamic miRNA–mRNA network identified several regulatory axes with relevance to placental and maternal vascular biology. *ADGRA2* was targeted by three independent miRNAs, miR-101-3p, miR-15a-5p, and miR-21-5p. *ADGRA2*, also known as *GPR124*, functions as a Wnt7 co-activator and has been implicated in vascular development, trophoblast invasion, and angiogenesis.^62,63^ The convergent targeting of *ADGRA2* suggests that it may represent a regulatory node within the temporally dynamic maternal blood network.

*HSD17B1* was also targeted by three miRNAs, miR-20a-5p, miR-106b-5p, and miR-3178. *HSD17B1* is involved in placental estrogen biosynthesis and has been implicated in trophoblast differentiation and PE-related placental dysfunction. ^64^ The miR-3178 → *HSD17B1* pair was the only non-canonical interaction in the stringent network, with both miRNA and target upregulated at M2. This exception highlights the complexity of miRNA-target relationships in vivo and suggests that some regulatory pairs may involve mechanisms beyond classical miRNA-mediated mRNA degradation, including indirect regulation, competing endogenous RNA effects, translational regulation, or post-transcriptional stabilization.

Additional prioritized pairs, including miR-15a-5p → ***KCP*** and miR-15a-5p → ***UHRF1***, point to TGF-β-related and epigenetic regulatory pathways. *KCP* regulates TGF-β/activin signaling through Smad2/3 inhibition,^65^ while UHRF1 is involved in epigenetic regulation through DNA methylation maintenance. These pathways are biologically plausible in PE given the central roles of trophoblast differentiation, invasion, immune tolerance, and developmental regulation in placental function.

Of the 29 miRNA–mRNA pairs identified, 8 showed direct or strong pathway-level convergence with recent placental studies implicating VGLL3, a Hippo pathway transcriptional coregulator, in PE-associated trophoblast dysfunction, endothelial pathology, immune dysregulation, and altered PI3K/AKT, FOXO, TGF-β, and trophoblast differentiation programs.^66^ In our maternal whole-blood data, the convergent targeting of ADGRA2 **(Wnt7 signaling)**, the miR-101-3p → DEPP1 axis **(PI3K/FOXO)**, HSD17B1-related regulatory pairs **(trophoblast differentiation)**, and the miR-15a-5p → KCP pair **(TGF-β pathway)** map onto several of these same biological programs. This cross-compartment convergence supports the hypothesis that a subset of maternal blood signatures may reflect placental or shared maternal–placental stress-response pathways. However, because our study did not directly profile PE placentas from the same pregnancies, these VGLL3–Hippo connections should be interpreted as mechanistic convergence rather than evidence of direct causal mediation.

A notable structural feature of the temporally dynamic network was the convergent targeting of several mRNAs by multiple independent miRNAs. ADGRA2 was targeted by three miRNAs — miR-101-3p, miR-15a-5p, and miR-21-5p — and HSD17B1 by three others — miR-20a-5p, miR-106b-5p, and miR-3178 — while additional targets including NRM and EEF2KMT were co-regulated by four or five miRNAs in the broader 29-pair network. Convergent multi-miRNA targeting of a common transcript is generally interpreted as evidence of buffered or reinforced regulation, in which the loss or attenuation of any single miRNA is insufficient to fully derepress the target, conferring robustness to the suppressive signal.^14,67^ In the context of temporal reversal observed here, convergent targeting may additionally reflect a sequential regulatory handoff, whereby different miRNAs — active at overlapping but distinct stages — collectively sustain target suppression across a broader gestational window than any single miRNA could achieve alone. Alternatively, independent miRNAs targeting the same transcript may contribute to fine-tuned magnitude control, enabling graded rather than binary regulation of target abundance in response to changing physiological states. The consistent co-targeting of *ADGRA2* and *HSD17B1* across multiple miRNAs in a temporally dynamic network therefore suggests that these transcripts represent high-priority regulatory nodes in PE-associated maternal blood, warranting prioritization in future functional and interventional studies.

### Progressive Heme-Metabolism-Related Transcriptional Suppression

In contrast to PI3K/AKT/mTORC1 signaling, heme metabolism showed progressive suppression across pregnancy, with negative enrichment observed at both early and late gestation and substantially stronger suppression at M2. This finding is distinct from the compensatory heme oxygenase-1 pathway, which has been widely studied in PE as an adaptive response to oxidative stress in placental tissue.^68–71^ Our data instead point to progressive suppression of heme-metabolism-related transcriptional programs in maternal whole blood. Importantly, erythroid and hemoglobin signature analyses did not support differential red blood cell contamination or hemolysis as an explanation for this signal.

The progressive transcriptional suppression of heme-metabolism programs observed here is contextually coherent with the known systemic iron and heme dysregulation of PE. Preeclampsia is associated with elevated maternal serum iron, ferritin, and transferrin saturation from the second trimester onward, alongside hemoconcentration due to impaired plasma volume expansion, and increased placental release of cell-free fetal hemoglobin into the maternal circulation — collectively representing a state of heme and iron excess rather than deficiency in the maternal circulation.^72–74^ In this context, transcriptional suppression of heme-biosynthetic and iron-regulatory programs in whole blood may reflect adaptive feedback responses to iron overload rather than primary heme deficiency, a distinction supported by the composition of the M2 leading edge genes identified in this study, which included the heme-sensing kinase EIF2AK1, the ferritinophagy receptor NCOA4, and the iron-sulfur cluster assembly factor ISCA1 — regulatory molecules whose transcriptional behavior is consistent with iron-replete stress states rather than erythroid depletion. Whether this transcriptional suppression represents a primary regulatory response, a compensatory adaptation, or a secondary consequence of altered erythroid or vascular biology in PE cannot be determined from the current data and warrants direct investigation using hematologic, proteomic, and functional assays in longitudinal PE cohorts.

### Placental Expression and Tissue-Compartment Relationships

Placental expression analyses provided biological support for selected blood-derived regulatory pairs. Most temporally dynamic miRNAs showed detectable placental expression in at least one resource, though miR-3178 showed absent or inconsistent placental expression across datasets, suggesting these pairs may reflect maternal systemic rather than placental regulatory programs. The strongest support was observed for miR-101-3p → *DEPP1*, ADGRA2-targeting pairs, and HSD17B1-targeting pairs. These findings suggest that at least a subset of maternal whole-blood signals may reflect placental or shared maternal–placental biology.

At the same time, placental concordance was incomplete, which is biologically informative. Some circulating signatures may derive from placental release into maternal blood through extracellular vesicles or cell-free RNA, as has been reported for selected PE-associated miRNAs.^75^ Others may reflect maternal systemic responses to placental dysfunction, including immune, vascular, endothelial, or metabolic adaptation. Therefore, the blood-derived signatures identified here should not be interpreted as exclusively placental in origin. Rather, they likely represent a mixture of placental, maternal systemic, and shared stress-response signals.

### Translational and Biomarker Implications

These findings have several implications for biomarker development. Most importantly, the lack of substantial same-direction temporal conservation suggests that PE biomarkers are likely to be stage-dependent rather than static. Early pregnancy signatures may be more relevant for risk prediction before symptom onset, whereas late pregnancy signatures may better reflect disease characterization, maternal response, or disease progression. This implies that validation studies should be carefully matched by gestational age and that biomarker panels may need to be timepoint-specific.

The distinction between dynamically remodeled pathways and progressively suppressed transcriptional programs may also help prioritize future mechanistic studies. Programs showing biphasicity, such as PI3K/AKT/mTORC1 signaling and the associated miRNA–mRNA network, may reflect stage-dependent regulatory adaptation. Programs showing progressive suppression, such as heme-metabolism-related transcriptional pathways, may reflect cumulative systemic stress or worsening metabolic dysregulation. Whether these pattern classes respond differently to intervention remains speculative and will require direct experimental testing.

Preclinical studies have suggested that modulation of PI3K/AKT/mTOR, heme oxygenase, and Hippo-related pathways can alter PE-relevant phenotypes in experimental systems.^51,66,68,71^ These studies provide biological context for the pathways identified here, but our findings should not be interpreted as therapeutic evidence. Any intervention-oriented interpretation will require direct testing in appropriate experimental models, with careful attention to gestational timing, dosing, safety, and pregnancy-specific physiology.

### Strengths and Limitations

This study has several strengths. The longitudinal design enabled direct comparison of molecular patterns across pregnancy rather than relying only on separate cross-sectional timepoints. The integration of miRNA and mRNA data from the same samples allowed evaluation of inverse regulation and network structure. Placental expression assessment across internal and external resources strengthened the biological interpretation of selected blood-derived signatures. Experimental validation of miR-101-3p-mediated DEPP1 repression provided functional support for one prioritized regulatory axis. Sensitivity analyses addressing maternal vitamin D status, estimated leukocyte composition, and disease severity supported the robustness of the main findings.

Several limitations should also be acknowledged. First, the study remains observational, and most network relationships are inferred from expression patterns and target databases rather than experimentally validated. Although the trophoblast experiments support miR-101-3p-mediated repression of DEPP1, they do not establish direct binding in vivo or prove that this interaction contributes causally to PE development. Second, whole-blood profiling integrates signals across multiple cell types and circulating compartments, limiting cell-type-specific inference. Although deconvolution analyses did not identify major leukocyte-composition differences, single-cell or cell-type-resolved profiling would provide more granular information. Third, placental expression support was incomplete and varied across pairs; although many selected miRNAs and target genes showed detectable placental expression in at least one resource, placental expression alone does not establish placental origin of circulating signatures. Fourth, the number of severe PE cases was limited, reducing power for severity-stratified analyses. Additionally, our primary differential expression and temporal classification analyses were conducted at the group level, and individual-level directional patterns were not systematically characterized. Whether the biphasic miRNA–mRNA patterns identified here are consistent across individual PE cases or reflect heterogeneity within the PE population — potentially related to disease severity, gestational age at onset, or other clinical features — remains to be determined. Single-patient-level analyses and clinically stratified longitudinal studies will be needed to assess the robustness and generalizability of the biphasic network architecture identified here. Fifth, downstream analyses focused primarily on protein-coding genes and miRNAs, while other regulatory layers, including long noncoding RNAs, were not deeply characterized. Finally, sampling windows within M1 and M2 covered several gestational weeks, and more granular sampling would be valuable for defining finer-scale temporal transitions.

### Conclusions

In summary, this longitudinal study shows that PE is associated with substantial temporal reorganization of maternal whole-blood miRNA–mRNA networks across pregnancy. Rather than showing stable molecular signatures, PE-associated expression patterns were predominantly stage-specific or biphasic. At the pathway level, PI3K/AKT/mTORC1 signaling showed dynamic remodeling, whereas heme-metabolism-related transcriptional programs showed progressive suppression. Integration of temporal miRNA–mRNA networks with placental expression and trophoblast validation prioritized several regulatory axes, including ADGRA2-targeting miRNAs, HSD17B1-targeting miRNAs, and the experimentally supported miR-101-3p → *DEPP1* interaction. These findings support a gestational-age-specific framework for PE biomarker development and identify temporally dynamic regulatory programs for future mechanistic investigation.

## Supporting information

Supplementary Methods, Results, Figures

Differential expression results are provided in Supplementary Data

GSEA results are provided in Supplementary Data 2

## Acknowledgements

We thank the participants and study staff of the VDAART study for their contributions to this research.

## Author contributions

HM designed, led the study and analysis and drafted the manuscript. STW was involved in trial design of VDAART. CJ and ZL, assisted in conduction of the data and statistical analyses. CJ and DH conducted experimental validations. STW, JL, CJ, DH, and ZL assisted in revision of the manuscript. All authors edited and approved the final manuscript.

## Data availability

Deidentified participant-level data from the Vitamin D Antenatal Asthma Reduction Trial (VDAART) may be made available to qualified investigators upon reasonable request and approval by the VDAART Data Coordinating Center, subject to applicable institutional, informed-consent, and data-use requirements. Requests should be directed to the corresponding author. Raw and processed mRNA-and miRNA-sequencing data generated in this study will be deposited in the NCBI Gene Expression Omnibus (GEO), and accession numbers will be provided before publication.

