## Supplementary Methods, Results, Figures for "Longitudinal miRNA-mRNA Profiling Reveals Network Reorganization in Preeclampsia"

**Longitudinal miRNA-mRNA Profiling Reveals Stage-Dependent Network Reorganization in Preeclampsia: Dynamic Pathway Reversal and Progressive Metabolic Suppression**

Mirzakhani *et al.*

**Supplementary materials in this file:**

**A. Supplementary Methods.**

1. VDAART Trial Eligibility and Exclusion Criteria

2. Vitamin D Measurement

3. Whole-Blood RNA Extraction, Library Preparation, and Sequencing

4. Sequencing Data Processing and Quality Control

5. Erythroid and Hemoglobin Signature Analysis

6. Placental RNA-seq Sample Processing and Expression Assessment

7. Public Placental Expression Databases

8. miRNA–mRNA Target Prediction and Network Integration

9. Cytoscape Network Construction and Hub Definitions

10. CIBERSORT Deconvolution and Cell-Composition Sensitivity Models

11. Experimental Validation Protocol: miR-101-3p Mimic Transfection, qPCR, and Western Blotting

12. Software Versions

**B. Supplementary Results**

1. RNA-seq QC, PCA, and erythroid/hemoglobin contamination assessment

2. Differential expression of noncoding and other annotated transcripts

3. Temporal classification of all M1-dysregulated mRNA-seq features

4. Expanded Hallmark pathway analysis and pathway-gene networks

5. Immunologic and cell-type GSEA

6. FOXO and PI3K-related supporting signatures

7. PI3K/AKT/mTORC1 pathway gene composition

8. Expanded miRNA–mRNA network analysis

9. Literature and mechanistic support for prioritized miRNA–mRNA pairs

10. Placental expression support for selected regulatory pairs

11. Sensitivity Analyses Confirm Robustness of Temporal Signatures.

**A. Supplementary Methods**

**Supplementary Methods 1. VDAART Trial Eligibility and Exclusion Criteria**

Key VDAART exclusion criteria included smoking or nicotine product use within 1 month of enrollment; chronic medical conditions, including diabetes mellitus and chronic hypertension requiring treatment; multiple gestation; assisted reproduction, including IUI or IVF; parathyroid or thyroid disease; nephrolithiasis; sarcoidosis; smoking, vitamin D supplementation exceeding 2,000 IU/day; and illicit drug use within 6 months of enrollment. Additional details of the trial design and primary findings have been reported previously.

**Supplementary Methods 2. Vitamin D Measurement**

Maternal 25-hydroxyvitamin D [25(OH)D] concentrations were measured at enrollment, 10–18 weeks’ gestation, and late pregnancy, 32–38 weeks’ gestation, using a chemiluminescence immunoassay, DiaSorin Liaison®. The assay detection limit was ≤4.0 ng/mL, with inter-assay coefficient of variation of 11.2% and intra-assay coefficient of variation of 8.1%. These measurements were used to describe baseline characteristics and in sensitivity analyses evaluating the potential influence of vitamin D status on differential expression results.

**Supplementary Methods 3. Whole-Blood RNA Extraction, Library Preparation, and Sequencing**

Total RNA was extracted from PAXgene Blood RNA tubes using the PAXgene Blood miRNA Kit, which permits simultaneous isolation of total RNA including small RNAs. RNA concentration was measured using the Qubit RNA HS Assay Kit, and RNA integrity was assessed using the Agilent 2100 Bioanalyzer with the RNA 6000 Nano Kit. Samples meeting prespecified quality thresholds, RIN ≥7.0 and RNA concentration ≥50 ng/μL for mRNA-seq or ≥25 ng/μL for miRNA-seq, were advanced to library preparation.

mRNA libraries were prepared from 450 ng total RNA using the TruSeq Stranded Total RNA Library Prep Kit with Ribo-Zero Globin. Libraries were quantified using the Qubit dsDNA HS Assay Kit, assessed on the Agilent 2100 Bioanalyzer using the High Sensitivity DNA Kit, normalized to equimolar concentrations, pooled, and sequenced on the Illumina NextSeq 550 platform using High Output flow cells, 150 cycles, to generate 75-bp paired-end reads. The target sequencing depth was 75 million reads per sample.

Small RNA libraries were prepared from 100 ng total RNA using the NEXTflex Small RNA Sequencing Kit v3 for Illumina Platforms. To reduce contamination from highly abundant hemolysis-associated miRNAs, hsa-miR-486-5p, hsa-miR-92a-3p, and hsa-miR-451a, sequence-specific erythrocyte miRNA blockers were added before adapter ligation. Libraries were quantified, normalized, pooled, and sequenced on the Illumina NextSeq 550 platform using High Output flow cells, 75 cycles, to generate single-end reads. The target sequencing depth was 12 million reads per sample.

**Supplementary Methods 4. Sequencing Data Processing and Quality Control**

Raw reads from mRNA-seq and miRNA-seq were processed using bcbio-nextgen version 1.2.9.59 For mRNA-seq, reads underwent quality assessment with FastQC version 0.11.9, adapter trimming and quality filtering with cutadapt version 3.4, alignment to GRCh38 with STAR version 2.7.10a, and gene-level quantification with featureCounts, Subread version 2.0.3, using GENCODE v46 annotations. Quality metrics included alignment rate, duplicate rate, ribosomal RNA content, and sequencing depth.

For miRNA-seq, reads underwent quality assessment with FastQC version 0.11.9, adapter trimming with cutadapt version 3.4, selection of reads 18–26 nucleotides in length, and alignment to miRBase release 22 with bowtie version 1.2.3. Quality metrics included adapter content, read-length distribution, alignment rate, and sequencing depth.

Samples were excluded if they had alignment rate <80% for mRNA-seq or <70% for miRNA-seq, or insufficient sequencing depth, defined as <20 million aligned reads for mRNA-seq or <5 million aligned reads for miRNA-seq.

Genes located on the X and Y chromosomes were excluded from differential expression analyses for two complementary reasons. First, fetal cell-free RNA circulating in maternal whole blood contains Y-linked transcripts in pregnancies with male fetuses, and differential fetal sex distribution between PE cases and controls could introduce spurious associations unrelated to maternal disease biology. Second, inter-individual variability in X-inactivation escape may contribute transcriptional noise that is not disease-related. Excluding sex chromosome genes addresses these sources of potential confounding at the feature level. Additionally, fetal sex was included as a covariate in all primary differential expression models to account for residual fetal sex-associated transcriptome differences in autosomal genes — including immune, vascular, and growth-related pathways known to differ by fetal sex in maternal circulation — that would not be captured by sex chromosome gene removal alone. Together, these two complementary adjustments ensure that neither sex chromosome gene expression nor broader fetal sex-associated transcriptome variation confounds the PE case-control comparison.

**Supplementary Methods 5. Erythroid and Hemoglobin Signature Analysis**

To evaluate whether heme-related findings could reflect erythroid contamination or hemolysis, we defined a 20-gene erythroid marker signature and a 9-gene hemoglobin signature. The erythroid signature included ALAS2, AHSP, SLC4A1, GYPA, GYPB, EPB42, ANK1, RHAG, BPGM, FECH, KLF1, HEMGN, CA1, CA2, SPTA1, SPTB, EPB41, GATA1, RHD, and RHCE. The hemoglobin signature included HBA1, HBA2, HBB, HBD, HBE1, HBG1, HBG2, HBZ, and HBM (**Supplementary Tables S1 and S2**).

Single-sample enrichment scores were calculated using the GSVA R package version 2.0.7 with ssGSEA applied to variance-stabilized gene expression values. The GSVA (Gene Set Variation Analysis) is a non-parametric, unsupervised method used to estimate variation in gene set enrichment over a sample population. Associations between PE status and each signature score were tested using logistic regression models adjusted for gestational age, maternal body mass index, parity, treatment arm, and race/ethnicity. Models were fit separately at M1 and M2, and two-sided P values <0.05 were considered statistically significant.

**Supplementary Methods 6. Placental RNA-seq Sample Processing and Expression Assessment**

Placental expression support was assessed using VDAART placental tissue collected from uncomplicated term pregnancies without preeclampsia, gestational diabetes, preterm birth, or other pregnancy complications. For placental miRNA-seq, villous tissue from 42 placentas was snap-frozen and stored at −80°C. Total RNA including small RNAs was extracted using the miRNeasy Mini Kit, and libraries were prepared using the NEXTflex Small RNA Sequencing Kit v3 from 100 ng total RNA. Libraries were sequenced on the Illumina NextSeq 550 platform to generate 75-bp single-end reads, targeting 12 million reads per sample.

For placental mRNA-seq, villous tissue from 28 placentas was fixed in 10% neutral-buffered formalin and embedded in paraffin. Total RNA was extracted from 5-μm FFPE sections using the AllPrep DNA/RNA FFPE Kit. Libraries were prepared from 50 ng total RNA using the SeqPlex RNA Amplification Kit and sequenced on the Illumina HiSeq 3000 platform to generate 50-bp single-end reads.^6^ Placental miRNA-seq data were processed using the same bcbio-nextgen workflow as whole-blood miRNA-seq. Placental FFPE mRNA-seq reads were aligned to GRCh38 and quantified at the gene level using featureCounts. Expression values were analyzed using DESeq2 with variance-stabilizing transformation. A feature was considered expressed in placenta if it had median log2 normalized count >5 across placental samples or was detected at ≥1 count in all samples. Placental expression was summarized using median and interquartile range of variance-stabilized counts.

**Supplementary Methods 7. Public Placental Expression Databases**

To complement internal placental RNA-seq data, public expression resources were queried. For mRNAs, the Human Protein Atlas, version 25.0, accessed March 10, 2026, was used to assess expression in normal placental tissue. Expression was categorized according to HPA annotations as Not detected, Low, Medium, or High. For miRNAs, the miRNA Tissue Expression Database, miTED, accessed March 10, 2026, was queried to assess placental expression. Expression levels were recorded as reads per million or normalized counts when available. Evidence from internal VDAART placental RNA-seq, HPA, and miTED was compiled for selected temporally dynamic miRNAs and target mRNAs.

**Supplementary Methods 8. miRNA–mRNA Target Prediction and Network Integration**

Putative regulatory relationships between differentially expressed miRNAs and mRNAs were identified using multiMiR version 1.32.0.65 Queried validated databases included miRTarBase, TarBase, and miRecords. Queried predicted databases included TargetScan, miRanda, DIANA-microT, EIMMo, MicroCosm, miRDB, and PicTar. Predicted interactions were restricted to those with scores above the 90th percentile where applicable.

For the early-pregnancy network, each differentially expressed miRNA was matched to predicted targets that were also differentially expressed at M1. miRNA–mRNA pairs were retained if the miRNA and mRNA showed inverse differential expression and if the interaction was supported by at least two independent databases or by at least one experimentally validated database. Retained pairs were then evaluated at M2 to determine whether both members remained dysregulated, reversed direction, or became M1-specific.

**Supplementary Methods 9. Cytoscape Network Construction and Hub Definitions**

Networks were visualized and analyzed in Cytoscape version 3.10.4. Hub miRNAs were defined as miRNAs targeting at least four differentially expressed mRNAs, and hub mRNAs were defined as transcripts targeted by at least two differentially expressed miRNAs. Hub analyses were performed separately for the broader M1 network, temporally filtered network, and stringent directional-reversal network.

**Supplementary Methods 10. CIBERSORT Deconvolution and Cell-Composition Sensitivity Models**

Relative leukocyte fractions were estimated using the CIBERSORT algorithm through the immunedeconv R package version 2.0.3.69, CIBERSORT used support vector regression and the LM22 signature matrix to estimate the relative abundance of 22 immune cell subtypes. Deconvolution was performed separately for M1 and M2 using variance-stabilized expression values with 1,000 permutations.

To reduce dimensionality, principal component analysis was performed on estimated leukocyte fractions, and the first three principal components were included as additional covariates in sensitivity models. Differential expression results from the primary and cell-composition-adjusted models were compared using Spearman correlation of log2 fold changes and overlap of significant features. Group differences in each estimated cell type were evaluated using Wilcoxon rank-sum tests separately at M1 and M2 with Benjamini–Hochberg correction.

**Supplementary Methods 11. Experimental Validation Protocol: miR-101-3p Mimic Transfection, qPCR, and Western Blotting**

HTR-8/SVneo trophoblast cells were transfected with hsa-miR-101-3p mimic or negative control, harvested at 24 and 48 hours post-transfection, and analyzed for miR-101-3p expression (qPCR), DEPP1 mRNA (qPCR), and DEPP1 protein (Western blot). Three independent experiments were performed with technical replicates.

HTR-8/SVneo cells, ATCC CRL-3271, were maintained in RPMI-1640 medium supplemented with 5% fetal bovine serum at 37°C in a humidified atmosphere containing 5% CO₂. Cells from passages 5–15 were plated in 6-well plates at 2.75 × 10⁵ cells/well and transfected at approximately 50–60% confluence with hsa-miR-101-3p mimic or negative control mimic at a final concentration of 20 nM using Lipofectamine 2000 in Opti-MEM. Cells were harvested at 24 and 48 hours after transfection. Three independent experiments were performed, with triplicate wells used for RNA analysis and duplicate wells used for protein analysis for each condition and timepoint.

For RNA analysis, total RNA including small RNAs was extracted using the miRNeasy Advanced Mini Kit. miRNA cDNA was synthesized from 12 ng total RNA using the TaqMan MicroRNA Reverse Transcription Kit and amplified using assays for hsa-miR-101-3p and RNU48. mRNA cDNA was synthesized from 500 ng total RNA using the High-Capacity cDNA Reverse Transcription Kit and amplified using assays for DEPP1 and ACTB. Quantitative PCR was performed using TaqMan Universal PCR Master Mix on an ABI PRISM 7900HT system. Each sample was assayed in technical triplicate.

For protein analysis, whole-cell lysates were prepared in RIPA buffer supplemented with protease and phosphatase inhibitors. Protein concentration was measured by DC Protein Assay. Equal amounts of protein were resolved on 4–15% Mini-PROTEAN TGX gels, transferred to PVDF membranes, and immunoblotted with antibodies against DEPP1 and α-actinin, followed by HRP-conjugated anti-rabbit IgG secondary antibody. Signals were detected by chemiluminescence and quantified with Image Lab. DEPP1 protein abundance was normalized to α-actinin for each sample.

**Supplementary Methods 12. Software Versions**

All statistical analyses were performed in R version 4.5.0. Key packages included DESeq2 version 1.50.2, multiMiR version 1.32.0, GSVA version 2.0.7, clusterProfiler version 4.18.4, immunedeconv version 2.0.3, and ggplot2 version 4.0.1. Cytoscape version 3.10.4 was used for network analysis and GraphPad Prism version 10.6.1 was used for experimental validation analyses.

**B. Supplementary Results**

**Supplementary Results 1. RNA-seq QC, PCA, and erythroid/hemoglobin contamination assessment**

Principal component analysis of variance-stabilized expression data showed substantial overlap between PE cases and controls at both timepoints. PC1 and PC2 captured 66% of variance for mRNA expression and 42% of variance for miRNA expression, indicating that PE status was not the dominant source of global expression variability.

Because Hallmark GSEA identified heme-metabolism dysregulation, we evaluated whether this signal could reflect differential red blood cell contamination or hemolysis. We calculated single-sample enrichment scores for a 20-gene erythroid marker signature and a 9-gene hemoglobin gene signature. Neither erythroid scores nor hemoglobin gene scores differed significantly between PE cases and controls at M1 or M2. Logistic regression models adjusted for gestational age, maternal body mass index, parity, treatment arm, and race/ethnicity did not identify significant associations between PE status and either erythroid or hemoglobin signatures. These analyses support the interpretation that heme-metabolism pathway suppression was not driven by differential erythroid contamination or hemolysis (**Supplementary Table S3**).

**Supplementary Results 2. Differential expression of noncoding and other annotated transcripts**

At early pregnancy, differential expression analysis identified 238 mRNA-seq features associated with subsequent PE, including 66 protein-coding genes, 120 long noncoding RNAs. Although the main analyses focused on protein-coding transcripts to support pathway enrichment and miRNA-target integration, noncoding and other annotated features accounted for the majority of early pregnancy differential expression. Directional asymmetry was observed across the full feature set, with 229 of 238 features downregulated and 9 upregulated in PE cases relative to controls.

These results indicate that early pregnancy PE-associated transcriptomic dysregulation extends beyond protein-coding genes and includes multiple classes of regulatory and noncoding transcripts. Full differential expression results for protein-coding and noncoding features at M1 and M2 are provided in **Supplementary Data 1**.

**Supplementary Results 3. Temporal classification of all M1-dysregulated mRNA-seq features**

To assess whether temporal reversal was restricted to protein-coding genes or reflected a broader transcriptomic pattern, we extended the temporal classification analysis to all 238 M1-dysregulated mRNA-seq features. In total, 78 features showed directional reversal at M2, whereas 160 features were M1-specific. Only 18 of the 78 directionally reversing features were protein-coding genes; the remainder included long noncoding RNAs, pseudogenes, and other annotated transcripts.

Effect sizes across all M1-dysregulated features showed a strong negative correlation between M1 and M2, consistent with coordinated temporal inversion rather than random fluctuation. This broader analysis supports the conclusion that PE-associated whole-blood transcriptional signatures are predominantly stage-dependent or directionally reversed rather than stably conserved across pregnancy. Full temporal classifications are provided in **Supplementary Data 1**.

**Supplementary Results 4. Expanded Hallmark pathway analysis and pathway-gene networks**

Hallmark GSEA identified distinct pathway profiles at M1 and M2. At M1, 15 pathways were significantly enriched with mixed directionality (4 upregulated, 11 downregulated), including positive enrichment of protein secretion, androgen response, UV response downregulated genes, and PI3K/AKT/mTORC1 signaling, together with negative enrichment of heme metabolism, cholesterol homeostasis, xenobiotic metabolism, interferon alpha response, estrogen response, DNA repair, E2F targets, G2M checkpoint, and MYC targets. The positive enrichment of the UV response downregulated gene set at M1 — representing genes normally suppressed following UV exposure — may reflect shared transcriptional regulatory programs with oxidative stress or DNA damage response pathways rather than literal UV-related biology, and is noted here for completeness without specific mechanistic interpretation. At M2, enriched pathways were uniformly downregulated, with strongest suppression of heme metabolism and reactive oxygen species pathway, followed by IL6/JAK/STAT3 signaling, PI3K/AKT/mTORC1 signaling, complement/interferon-related pathways, and programmed cell-death-related pathways.

Pathway-gene network visualization (**Figure 2C-D**) showed broader and more heterogeneous pathway architecture at M1, with both positively and negatively enriched modules. In contrast, the M2 network was more restricted and uniformly downregulated, with heme metabolism forming the dominant suppressed module. These network patterns were consistent with the main GSEA results and supported the interpretation that PE-associated pathway structure changes substantially across gestation. Complete Hallmark GSEA results and leading-edge genes are provided in **Supplementary Table S4**.

**Supplementary Results 5. Immunologic and cell-type GSEA**

To further characterize biological programs underlying early pregnancy dysregulation, we performed GSEA using MSigDB immunologic signatures, C7, and cell-type signatures, C8. At M1, 1,604 of 5,205 C7 immunologic gene sets were significantly enriched at FDR < 0.05, including immune cell activation states, vaccine-response signatures, and T-cell and B-cell differentiation programs. Among C8 cell-type signatures, 160 of 864 gene sets were significantly enriched at M1, including erythroid, NK/NKT, endothelial, and other cell-type-associated signatures.

In contrast, no C7 or C8 gene sets reached FDR < 0.05 at M2. This marked difference between M1 and M2 suggests that immune and cell-type-associated transcriptional signatures were primarily early-stage features, whereas Hallmark metabolic and signaling pathways showed more persistent or progressive dysregulation across pregnancy. Complete C7 and C8 GSEA results are provided in **Supplementary Data 2**.

**Supplementary Results 6. FOXO and PI3K-related supporting signatures**

FOXO-related immunologic signatures provided additional support for the PI3K/AKT–FOXO interpretation. At M1, FOXO target-related signatures showed bidirectional enrichment consistent with FOXO transcriptional inactivation during PI3K/AKT activation. Specifically, gene sets representing genes normally suppressed by FOXO were positively enriched (NES=+2.37), whereas FOXO target gene sets were negatively enriched (NES=-2.06). PI3K inhibitor-related signatures showed parallel support for PI3K pathway activation at M1.

DEPP1 appeared in multiple vascular endothelial cell-type signatures that were downregulated at M1, providing additional support for its relevance to vascular or endothelial-associated regulatory programs. Together, these analyses support the connection between M1 PI3K/AKT/mTORC1 activation, FOXO target suppression, and DEPP1 downregulation. Full FOXO, PI3K inhibitor, and related C7/C8 signature results are provided in **Supplementary Data 2**.

**Supplementary Results 7. PI3K/AKT/mTORC1 pathway gene composition**

To examine whether PI3K/AKT/mTORC1 pathway reversal reflected uniform regulation of the same genes or remodeling of pathway composition, we evaluated leading-edge genes contributing to pathway enrichment at M1 and M2. The M1-enriched pathway included upstream and regulatory components such as PTEN, GSK3B, SMAD2, and MAPK1, whereas M2-enriched pathway components included AKT1, GRB2, TSC2, MYD88, and TRIB3. Network visualization showed spatial separation of M1- and M2-enriched pathway components, with central nodes connecting stage-specific modules (**Figure 3A–B**).

Integration of temporally dynamic miRNAs with the PI3K/AKT/mTORC1 pathway network showed that multiple reversing miRNAs were predicted to target pathway components, including central or highly connected nodes. These findings suggest that miRNA-mediated regulation may contribute to pathway-level temporal remodeling. Detailed pathway genes and leading-edge membership are provided in **Supplementary Table S4**.

**Supplementary Results 8. Expanded miRNA–mRNA network analysis**

We constructed miRNA–mRNA regulatory networks at three levels of temporal stringency. This progressive refinement strategy reduced network complexity while enriching for temporally coherent regulatory interactions, **ultimately** yielding 29 high-confidence pairs at maximum stringency.

First, all 65 M1-dysregulated miRNAs were integrated with the 66 M1-dysregulated protein-coding genes to identify early pregnancy inverse regulatory pairs. This broad M1 network included key PE-relevant targets such as HSD17B1, DEPP1, UHRF1, MMRN2, ADGRA2, and CD109, each targeted by one or more differentially expressed miRNAs.

Second, we restricted the network to the 11 miRNAs showing temporal dysregulation across M1 and M2. This intermediate network retained a focused subset of target genes, including HSD17B1, DEPP1, ADGRA2, NRM, EEF2KMT, and PHKG1. Third, the most stringent network integrated the 11 temporally dynamic miRNAs with 18 directionally reversing protein-coding genes, yielding 29 high-confidence miRNA–mRNA pairs involving 9 miRNAs and 14 target genes.

The stringent network showed evidence of combinatorial regulation. miR-15a-5p was the most connected miRNA, targeting 7 genes, followed by miR-106b-5p, miR-101-3p, and miR-20a-5p. Several target genes were co-regulated by multiple miRNAs, including NRM, EEF2KMT, ADGRA2, HSD17B1, and PHKG1. These patterns support the presence of multi-miRNA regulatory architecture rather than isolated one-to-one miRNA-target relationships. Full predicted and validated target evidence for all 29 pairs is provided in **Supplementary Table S5**.

**Supplementary Results 9. Literature and mechanistic support for prioritized miRNA–mRNA pairs**

Among the 29 high-confidence temporally dynamic miRNA–mRNA pairs, several were prioritized based on temporal reversal, convergent miRNA targeting, placental expression, and literature support for relevance to PE or placental biology. ADGRA2 was targeted by miR-101-3p, miR-15a-5p, and miR-21-5p and has been implicated in Wnt7 signaling, vascular development, trophoblast invasion, and angiogenesis. HSD17B1 was targeted by miR-20a-5p, miR-106b-5p, and miR-3178 and is involved in placental estrogen biosynthesis and trophoblast differentiation. DEPP1 was targeted by miR-101-3p and linked the temporal miRNA–mRNA network to FOXO-mediated stress-response and autophagy-related programs.

Additional prioritized pairs included miR-15a-5p → *KCP*, linking to TGF-β pathway regulation; miR-15a-5p → *UHRF1,* linking to epigenetic regulation; miR-26b-5p → *APOA2*, linking to lipid metabolism; and miR-19b-3p → *EEF2KMT*, linking to translational control. Detailed PubMed-supported literature evidence, placental-expression annotation, and mechanistic comments for each pair are provided in **Table 2** and **Supplementary Table S6**.

**Supplementary Results 10. Placental expression support for selected regulatory pairs**

We assessed placental expression support for temporally dynamic miRNAs and their target genes using internal VDAART placental RNA-seq data and public expression resources. Most temporally dynamic miRNAs showed detectable placental expression in at least one expression resource, although expression levels varied across datasets. **miR-21-5p**, **miR-26b-5p**, and **miR-101-3p** showed the strongest and most consistent placental expression across datasets. In contrast, **miR-3178** and **miR-454-3p** showed weak, inconsistent, or absent placental expression, suggesting that regulatory pairs involving these miRNAs may be more reflective of maternal blood or systemic responses than placental expression programs.

Placental mRNA expression was also evaluated for selected directionally reversing protein-coding genes considered during miRNA–mRNA prioritization, including genes retained in the final high-confidence network and genes not retained after target-prediction filtering. **C1orf54**, although differentially expressed and included in the placental expression assessment, was not retained in the final 29-pair network because it did not meet high-confidence miRNA-target prediction criteria. Among target genes retained in the final temporally dynamic miRNA–mRNA network, placental expression support was strongest for **HSD17B1**, **DEPP1**, and **ADGRA2**.

Overall, these data support the biological plausibility of selected blood-derived regulatory pairs, particularly **miR-101-3p → DEPP1**, **ADGRA2**-targeting pairs, and **HSD17B1**-targeting pairs, while indicating that placental support is incomplete across the full network. Placental expression distributions are shown in **Supplementary Figure S1**, and pair-level placental evidence is summarized in **Supplementary Table S6**.

**Supplementary Results 11. Sensitivity Analyses Confirm Robustness of Temporal Signatures.** To assess whether identified temporal signatures were confounded by disease severity, cellular composition, or maternal vitamin D status, we performed three sensitivity analyses:

**Preeclampsia severity does not affect temporal gene signatures.** To assess whether temporally dynamic miRNA-mRNA signatures differed by disease severity, we compared gene expression between women with severe versus mild/moderate preeclampsia. None of the 18 directionally reversing protein-coding genes or 10 directionally reversing miRNAs showed differential expression between severe and mild PE cases (all FDR > 0.05). Similarly, no differences were observed for the 48 M1-specific genes or 54 M1-specific miRNAs. This indicates that temporal regulatory signatures identified in this study are shared across the PE severity spectrum and likely reflect common molecular mechanisms underlying PE pathogenesis, rather than severity-dependent pathways. The lack of severity-associated differences strengthens the generalizability of our findings to the broader PE population.

**Cell composition does not confound primary findings.** Computational cell type deconvolution (CIBERSORT, LM22 signature) revealed no significant differences in major leukocyte populations between PE cases and controls at either timepoint (**Supplementary Tables: Table S7**). Neutrophils (M1: P=0.119, M2: P=0.944), monocytes (M1: P=0.067, M2: P=0.372), T cells, and NK cells all showed similar proportions between groups. Only T cell gamma delta showed nominally significant difference at M1 (P=0.047) but did not survive multiple testing correction (Bonferroni-adjusted P > 0.05 for 22 comparisons). As a sensitivity analysis, we included cell composition principal components (capturing 99% of variance at M2; 98% at M1) as covariates in differential expression models. Effect sizes remained highly correlated between primary and cell composition-adjusted models (M1: r=0.87, P<0.0001; M2: r=0.90, P<0.0001), confirming that observed transcriptional differences reflect molecular dysregulation within cell types rather than compositional shifts. Given the absence of compositional differences and high effect size concordance, primary model results are presented throughout this manuscript.

**Maternal vitamin D status does not modify temporal gene signatures.** Maternal vitamin D status differed between PE cases and controls at early pregnancy (M1: 19.56±8.41 ng/mL vs. 24.88±14.56 ng/mL, P=0.026) but not at late pregnancy (M2: 30.09±15.28 ng/mL vs. 34.39±17.44 ng/mL, P=0.161). Because PE cases also had higher BMI at enrollment, and BMI was included in the primary differential expression models, we evaluated the correlation between BMI and 25(OH)D. Maternal BMI at enrollment was modestly but significantly inversely correlated with baseline 25(OH)D concentration (Spearman ρ = −0.32, P = 3.44 × 10⁻¹⁴; Pearson r = −0.29, P = 5.46 × 10⁻¹²; N = 768). To assess whether the baseline vitamin D difference influenced our findings, we repeated differential expression analyses with serum 25(OH)D concentration included as an additional covariate in DESeq2 models. Effect size estimates remained highly concordant between primary and vitamin D-adjusted models (Spearman correlation r ≥ 0.90, P < 0.0001 at both timepoints), and none of the temporally dynamic genes (18 directional reversals, 43 M1-specific protein-coding genes, 10 directional reversal miRNAs) showed substantial changes in significance or effect size after adjustment. This demonstrates that identified temporal regulatory signatures are robust to maternal vitamin D status and reflect PE pathophysiology independent of vitamin D concentrations, despite the baseline vitamin D difference between groups.

**C. Supplementary Figure:**

| **A.** | **B.** |
| --- | --- |
| 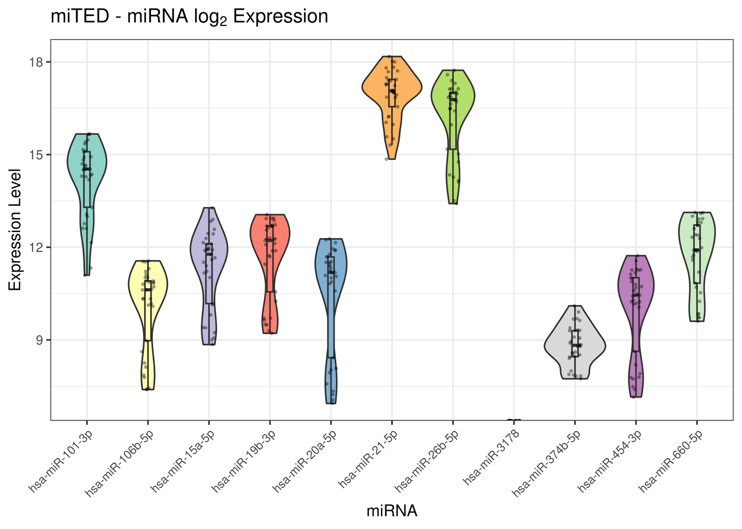 | 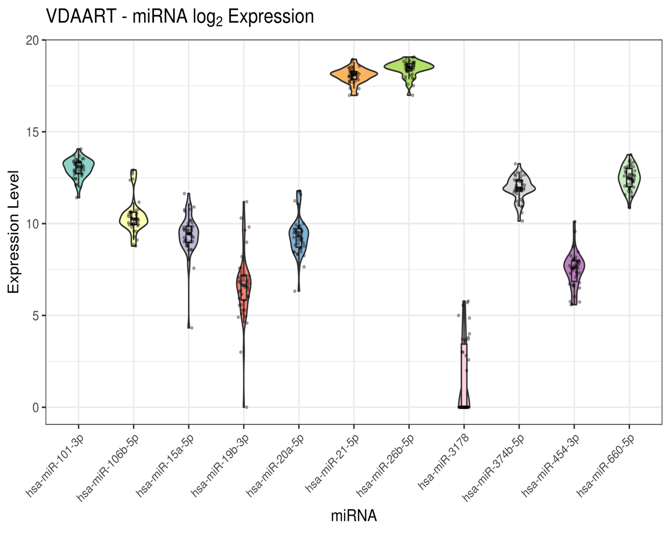 |
| **C.** | |
| 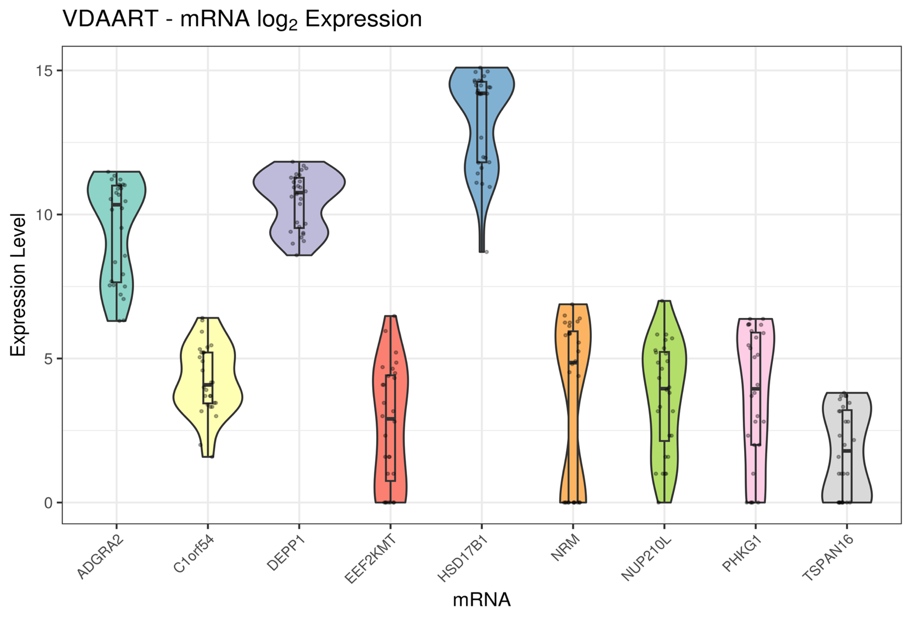 | |

**Figure S1. Placental expression support for temporally dynamic miRNAs and selected directionally reversing target genes.**

**(A)** miTED database miRNA expression levels (log₂ normalized counts) for 11 miRNAs in human placental tissue. Violin plots show expression distribution across multiple placental samples. miR-21-5p and miR-26b-5p show very high, consistent expression (log₂ ≈ 17-18); miR-101-3p shows high expression (log₂ ≈ 15); miR-3178 shows very low expression.

**(B)** VDAART cohort placental miRNA expression (log₂ normalized counts). Violin width indicates expression distribution across individual samples. Note excellent concordance with miTED for miR-21-5p, miR-26b-5p, and miR-101-3p (narrow distributions, high expression). miR-15a-5p, miR-19b-3p, and miR-3178 show wide distributions indicating high inter-sample variability or low/absent expression in many samples.

**(C)** VDAART placental mRNA expression, log₂ normalized counts, for selected directionally reversing protein-coding genes evaluated during miRNA–mRNA prioritization. **HSD17B1** showed very high expression with a narrow distribution, while **DEPP1** and **ADGRA2** showed robust expression. Other genes, including **EEF2KMT**, **NRM**, **NUP210L**, **PHKG1**, **C1orf54**, and **TSPAN16**, showed lower or more variable expression. **C1orf54** was differentially expressed and evaluated for placental expression support but was not retained in the final high-confidence predicted miRNA–mRNA network because it did not meet target-prediction confidence criteria.
