## Supplementary Tables S1-S8 for "Longitudinal miRNA-mRNA Profiling Reveals Network Reorganization in Preeclampsia"

**Table S1.** Study subjects’ characteristics

|  | PE cases  (N = 43) | Healthy  (N = 106) | P value |
| --- | --- | --- | --- |
| **Demographic** |  |  |  |
| Race, N (%)  White  Black/African American  Other/multiple | 15 (34.9)  21 (48.8)  7 (16.3) | 42 (39.6)  57 (53.8)  7 (6.6) | 0.185 |
| Treatment arm (4400 IU/day), N (%) | 22 (51.2) | 48 (45.3) | 0.638 |
| Clinical Enrollment Site, N (%)  Boston, MA  San Diego, CA  St. Louis, MO | 10 (23.3)  14 (32.6)  19 (44.2) | 17 (16.0)  34 (32.1)  55 (51.9) | 0.536 |
| Maternal Age, yr, mean±SD | 25.73±5.14 | 26.70±5.13 | 0.298 |
| Maternal Education, N (%)  Less than high school  Less than college  College or more | 10 (23.3)  16 (37.2)  17 (39.5) | 15 (14.2)  35 (33.0)  56 (52.8) | 0.250 |
| Income, N (%)  Less than 50k  More than 50k  Unknown | 16 (37.2)  11 (25.6)  14 (32.6) | 55 (51.9)  24 (22.6)  19 (17.9) | 0.184 |
| Income, N (%)  Less than 75k  More than 75k  Unknown | 21 (48.8)  6 (14.0)  14 (32.6) | 67 (63.2)  12 (11.3)  19 (17.9) | 0.205 |
| Marital Status, N (%)  Married  Divorced/separated  Not married | 16 (37.2)  26 (60.5)  1 (2.3) | 38 (35.8)  63 (59.4)  5 (4.7) | 0.796 |
| Parity, N (%)  More than 1^st^ pregnancy | 24 (55.8) | 68 (64.2) | 0.446 |
| Baseline 25(OH)D, ng/mL, mean±SD | 19.56±8.41 | 24.88±14.56 | 0.026 |
| 2^nd^ Timepoint (32-38 wks) 25(OH)D, ng/mL, mean±SD | 30.09±15.28 | 34.39±17.44 | 0.161 |
| Pre-pregnancy BMI, kg/m^2,^ mean±SD | 29.47±7.21 | 26.80±7.02 | 0.038 |
| Maternal BMI at Enrollment, mean±SD | 31.00±8.17 | 27.79±7.69 | 0.025 |
| Gestational Age at Enrollment (10-18 wks), mean±SD | 13.84±2.74 | 14.20±2.71 | 0.472 |
| Gestational Age at 2^nd^ Timepoint (32-38 wks), mean±SD | 33.07±3.26 | 33.57±1.65 | 0.218 |
| Maternal Asthma, N (%) | 20 (46.5) | 45 (42.5) | 0.787 |
| Fetal Sex (Male), N (%) | 24 (55.8) | 51 (48.1) | 0.502 |
| Gestational Age at Delivery, wks, mean±SD | 37.74±3.39 | 39.15±1.02 | < 0.001 |
| **Features of Cases** |  |  |  |
| Severity, N (%) | 11 (25.6) | - | < 0.001 |
| Preterm, N (%)  Very Preterm (<34 wks)  Moderate-Late Preterm (34–37 wks)  Term (≥37 wks) | 5 (11.6)  6 (14.0)  32 (74.4) | 0 (0.0)  0 (0.0)  106 (100.0) | - |

**Table S2. Erythroid Cell Marker Signature (20 genes):**

| **Gene Symbol** | **Gene Name** | **Function** | **Rationale for Inclusion** |
| --- | --- | --- | --- |
| ALAS2 | δ-aminolevulinic acid synthase 2 | Rate-limiting enzyme in heme biosynthesis | Erythroid-specific isoform |
| AHSP | Alpha-hemoglobin stabilizing protein | Chaperone for free α-globin chains | RBC-specific expression |
| SLC4A1 | Solute carrier family 4 member 1 (Band 3) | Major RBC membrane anion exchanger | Abundant RBC membrane protein |
| GYPA | Glycophorin A | Major RBC surface glycoprotein | RBC surface marker |
| GYPB | Glycophorin B | RBC surface glycoprotein | RBC surface marker |
| EPB42 | Erythrocyte membrane protein band 4.2 | RBC membrane cytoskeleton component | RBC membrane integrity |
| ANK1 | Ankyrin 1 | RBC membrane-cytoskeleton linker | RBC structural protein |
| RHAG | Rh-associated glycoprotein | Ammonium transporter | RBC-specific function |
| BPGM | Bisphosphoglycerate mutase | Catalyzes 2,3-BPG synthesis | Regulates hemoglobin oxygen affinity |
| FECH | Ferrochelatase | Terminal enzyme in heme biosynthesis | Incorporates iron into protoporphyrin IX |
| KLF1 | Krüppel-like factor 1 | Erythroid-specific transcription factor | Master regulator of erythropoiesis |
| HEMGN | Hemogen | Erythroid differentiation regulator | Promotes erythroid maturation |
| CA1 | Carbonic anhydrase 1 | CO2 transport | RBC-specific isoform |
| CA2 | Carbonic anhydrase 2 | CO2 transport | RBC-enriched isoform |
| SPTA1 | Spectrin alpha chain, erythrocytic 1 | RBC membrane cytoskeleton | Maintains cell shape |
| SPTB | Spectrin beta chain, erythrocytic | RBC membrane cytoskeleton | Maintains cell deformability |
| EPB41 | Erythrocyte membrane protein band 4.1 | RBC membrane-cytoskeleton linker | Stabilizes membrane structure |
| GATA1 | GATA binding protein 1 | Master erythroid transcription factor | Essential for erythroid lineage |
| RHD | Rh blood group D antigen | Defines Rh+ phenotype | RBC surface antigen |
| RHCE | Rh blood group CE antigens | Rh blood group system | RBC surface antigens |

**Table S3. Hemoglobin Gene Signature (9 genes):**

| **Gene Symbol** | **Protein** | **Type** | **Expression Pattern** |
| --- | --- | --- | --- |
| HBA1 | α-globin 1 | Adult α-globin | Major adult hemoglobin component |
| HBA2 | α-globin 2 | Adult α-globin | Major adult hemoglobin component |
| HBB | β-globin | Adult β-globin | Forms HbA (α2β2) - major adult hemoglobin |
| HBD | δ-globin | Minor adult β-like globin | Forms HbA2 (α2δ2) - minor adult hemoglobin |
| HBE1 | ε-globin | Embryonic globin | Embryonic hemoglobin component |
| HBG1 | γ-globin G | Fetal globin | Forms HbF (α2γ2) - fetal hemoglobin |
| HBG2 | γ-globin A | Fetal globin | Forms HbF (α2γ2) - fetal hemoglobin |
| HBZ | ζ-globin | Embryonic α-like globin | Embryonic hemoglobin component |
| HBM | μ-globin | Embryonic globin | Embryonic hemoglobin component |

**Table S4: RBC Contamination Quality Control**

| Signature | Timepoint | β (PE vs Control) | P-value | OR (95% CI) |
| --- | --- | --- | --- | --- |
| Erythroid markers (20 genes) | M1 | 0.10 (-0.63, 0.85) | 0.78 | 1.11 **(0.53, 2.34)** |
| Erythroid markers (20 genes) | M2 | -0.51 (-1.27, 0.23) | 0.18 | **0.60 (0.28, 1.26)** |
| Hemoglobin genes (9 genes) | M1 | -0.36 (-1.25, 0.51) | 0.42 | **0.70 (0.29, 1.67)** |
| Hemoglobin genes (9 genes) | M2 | -0.71 (-1.62, 0.15) | 0.11 | 0.49 **(0.20, 1.16)** |

**Table S5.** Enriched pathways at early (M1:10-18 weeks) and late pregnancy (32-38 weeks) and pathway genes.

| **Timepoint** | **Pathway** | **Description /**  **Category** | **Size** | **Leading Edge Number** | **NES** | **FDR** | **Direction** | **Pathway Genes** |
| --- | --- | --- | --- | --- | --- | --- | --- | --- |
| M1 | UV RESPONSE DN | UV response: downregulated genes | 141 | 75 | 2.015 | 0.024 | Up | DAB2, LPAR1, TFPI, SCAF8, CDON, TGFBR2, ATXN1, MAP2K5, TENT4A, PRDM2, PDLIM5, ATP2B4, PRKCA, NR3C1, PTPN21, CELF2, ATP2B1, RASA2, MAPK14, RUNX1, PTEN, AKT3, CDK13, KALRN, KIT, DYRK1A, NOTCH2, NIPBL, IGF1R, PLPP3, NEK7, SIPA1L1, SYNE1, BMPR1A, INPP4B, SPOP, INSIG1, SCN8A, PTPRM, ADD3, TGFBR3, BCKDHB, DLG1, DLC1, ADORA2B, NR1D2, MAGI2, FYN, SFMBT1, COL11A1, ATP2C1, ANXA4, YTHDC1, PRKAR2B, PMP22, SRI, LTBP1, PHF3, ADGRL2, GCNT1, NFKB1, MRPS31, NFIB, PPARG, MAP1B, NRP1, COL1A1, PRKCE, ICA1, VLDLR, AGGF1, RND3, HAS2, ITGB3, CDKN1B |
| M1 | PROTEIN SECRETION | protein secretion | 88 | 54 | 1.976 | 0.024 | Up | AP1G1, RAB22A, ARFIP1, STX16, ARFGEF1, MON2, SEC22B, GALC, STAM, SNAP23, KIF1B, STX12, RAB14, ARCN1, ARFGAP3, VPS4B, SGMS1, CLTC, SEC31A, RAB5A, DST, ADAM10, ARFGEF2, STX7, VPS45, ATP6V1H, RAB2A, VAMP4, VAMP3, MAPK1, COG2, CTSC, COPB1, SCAMP1, CAV2, BET1, USO1, AP3B1, LMAN1, TMED10, COPB2, GOLGA4, PPT1, DOP1A, TPD52, TSG101, ICA1, ZW10, SNX2, TMED2, EGFR, TMX1, ABCA1, CLCN3 |
| M1 | ANDROGEN RESPONSE | androgen response | 97 | 37 | 1.621 | 0.024 | Up | SGK1, TSC22D1, PDLIM5, CDK6, STK39, NGLY1, PTPN21, ELK4, GPD1L, GNAI3, RAB4A, GUCY1A1, B4GALT1, STEAP4, SRP19, UBE2J1, ELOVL5, SPCS3, VAPA, HMGCR, INPP4B, INSIG1, APPBP2, IQGAP2, CDC14B, SLC38A2, PIAS1, HMGCS1, SLC26A2, ABHD2, ACSL3, MAK, TMEM50A, MAF, LMAN1, TNFAIP8, ITGAV |
| M1 | PI3K AKT MTOR SIGNALING | PI3K signaling via AKT to mTORC1 | 103 | 34 | 1.457 | 0.028 | Up | GSK3B, PDK1, RIT1, THEM4, PIKFYVE, MAPK9, SMAD2, RAC1, VAV3, ITPR2, YWHAB, IRAK4, PRKAR2A, PTEN, MAPK8, PLA2G12A, CLTC, PRKCB, MAP3K7, ACTR3, ACTR2, PPP2R1B, CAB39, CAMK4, UBE2D3, ATF1, MAPK1, PLCB1, MAPK10, EIF4E, ARPC3, NCK1, RALB, SLA |
| M1 | ESTROGEN RESPONSE LATE | late estrogen response | 194 | 69 | -1.39 | 0.037 | Down | ANXA9, CDC20, KIF20A, NAB2, GINS2, SLC1A4, CAV1, SLC22A5, IDH2, IL17RB, RNASEH2A, FKBP4, IGFBP4, CACNA2D2, ABCA3, GALE, DCXR, SCNN1A, LSR, TOP2A, BLVRB, SLC7A5, MAPT, METTL3, GFUS, GPER1, TJP3, MICB, SEMA3B, RPS6KA2, KCNK5, SLC2A8, CKB, ALDH3B1, TMPRSS3, CXCL12, FABP5, SFN, BATF, ST14, ST6GALNAC2, DUSP2, GAL, AMFR, SLC29A1, PTPN6, TFF3, NHERF1, SIAH2, DLG5, SLC27A2, TST, WFS1, ISG20, XRCC3, DHCR7, PLXNB1, LLGL2, CLIC3, ITPK1, PAPSS2, KLF4, NCOR2, OPN3, CELSR2, NBL1, FLNB, HOMER2, TRIM29 |
| M1 | XENOBIOTIC METABOLISM | metabolism of xenobiotics | 196 | 55 | -1.429 | 0.028 | Down | TKFC, LCAT, ACO2, MTHFD1, ACP2, RETSAT, CYP1A1, IRF8, ESR1, ABHD6, KARS1, PYCR1, SHMT2, IGFBP4, EPHX1, PTGDS, ACOX3, DDT, PC, ACSM1, LONP1, DCXR, BLVRB, SLC12A4, DHRS1, TMEM97, HES6, DHPS, PSMB10, CFB, IGF1, COMT, ENTPD5, HMOX1, GSTM4, JUP, SERTAD1, LEAP2, BCAR1, CNDP2, AHCY, ANGPTL3, ASL, GSS, DDAH2, POR, PINK1, SMOX, NMT1, NPC1, ATOH8, FAH, MPP2, VTN, ECH1 |
| M1 | G2M CHECKPOINT | cell cycle progression: G2/M checkpoint | 192 | 65 | -1.446 | 0.028 | Down | NEK2, CDC20, MCM2, DDX39A, GINS2, CCNF, CDK4, ATF5, PRC1, KIF2C, HMGA1, CDC45, UCK2, CKS1B, UBE2C, LIG3, RAD54L, KIF22, HMMR, BIRC5, MCM3, CCNB2, CDKN2C, DTYMK, PTTG1, H2AX, TOP2A, TPX2, CHEK1, SLC7A5, TGFB1, CHAF1A, CENPF, UBE2S, CDK1, TFDP1, CCNA2, ESPL1, E2F2, RPA2, ABL1, CHMP1A, NASP, NUMA1, KPNA2, KIF11, E2F4, MCM5, CDC25B, MYC, BCL3, MYBL2, ILF3, BUB1, NSD2, FANCC, MKI67, POLE, POLQ, CDC25A, TNPO2, UPF1, CDKN3, CENPE, INCENP |
| M1 | MYOGENESIS | muscle differentiation | 195 | 90 | -1.525 | 0.024 | Down | MAPK12, CHRNB1, MYL6B, REEP1, DENND2B, PC, MYO1C, ITGA7, ATP2A1, SGCA, MYH11, PFKM, TGFB1, CRAT, SMTN, TAGLN, SH2B1, TPM2, LSP1, IGFBP7, DMPK, SPHK1, ACHE, SORBS3, AKT2, IGF1, PGAM2, KCNH2, CKB, PPFIA4, ITGB4, SIRT2, TCAP, GAA, SPEG, PLXNB2, CDH13, PICK1, PSEN2, BIN1, OCEL1, FLII, COL6A2, MYL4, CACNA1H, MYH9, ERBB3, TSC2, AGRN, FDPS, ACTA1, TNNT1, PYGM, SYNGR2, HDAC5, MYBPH, GSN, CKMT2, VIPR1, CTF1, EPHB3, GADD45B, DES, COL6A3, TNNT3, APP, CHRNG, ADCY9, MYH2, TNNI2, RYR1, ANKRD2, FGF2, FABP3, SCD, AK1, COL3A1, KIFC3, CFD, COX7A1, TNNC2, NOTCH1, NQO1, BAG1, CAMK2B, KCNH1, PDLIM7, AEBP1, MYLK, COL4A2 |
| M1 | CHOLESTEROL HOMEOSTASIS | cholesterol homeostasis | 71 | 35 | -1.543 | 0.028 | Down | ATF5, PCYT2, CXCL16, LSS, MVK, TMEM97, TM7SF2, GLDC, SEMA3B, PMVK, GSTM2, CHKA, TRIB3, FABP5, FBXO6, MVD, ACAT2, GUSB, ETHE1, CYP51A1, CBS, FASN, LGALS3, LGMN, FDPS, ECH1, JAG1, STX5, DHCR7, ABCA2, PLAUR, TNFRSF12A, SREBF2, ACTG1, LPL |
| M1 | HEME METABOLISM | heme metabolism and erythroid lineage | 187 | 84 | -1.56 | 0.024 | Down | HBD, HTRA2, HBB, ACP5, KHNYN, AHSP, H4C3, PRDX2, UROD, HBQ1, PPOX, GLRX5, PC, RHD, EPOR, RBM38, TSPO2, PIGQ, SPTB, OSBP2, H1-0, SLC2A1, BLVRB, C3, TMCC2, UBAC1, RAD23A, PDZK1IP1, ALAD, HMBS, E2F2, SIDT2, FTCD, HAGH, PPP2R5B, CTNS, BPGM, KLF1, AQP3, DCAF11, BSG, SELENBP1, CCND3, RHCE, TRIM10, MAP2K3, SLC66A2, FECH, MARCHF2, NFE2L1, ERMAP, SMOX, KEL, SLC4A1, RNF123, P4HA2, MYL4, HDGF, SLC6A9, DMTN, UROS, SLC22A4, GYPC, EPB42, FOXO3, CTSB, SLC25A38, TAL1, GYPB, SLC25A37, ANK1, LRP10, NUDT4, LMO2, MFHAS1, ALDH6A1, ADD2, NFE2, TNS1, UCP2, MXI1, MARCHF8, RCL1, CA1 |
| M1 | INTERFERON ALPHA RESPONSE | interferon alpha response | 97 | 53 | -1.606 | 0.028 | Down | TAP1, CNP, PSMB8, WARS1, IFITM3, BST2, IFI35, IRF7, CD74, PSMA3, CMTR1, RNF31, TRIM14, TRIM26, OGFR, HELZ2, ISG15, SLC25A28, IFITM1, GMPR, DHX58, BATF2, MOV10, TMEM140, PSME2, IFIT3, LAP3, LY6E, RSAD2, PSME1, UBA7, ISG20, LPAR6, UBE2L6, CMPK2, HLA-C, MX1, MVB12A, IFI30, IFIT2, PARP14, OASL, STAT2, LAMP3, LGALS3BP, PSMB9, CXCL10, OAS1, SAMD9, IRF1, TRIM21, IFITM2, IRF9 |
| M1 | UV RESPONSE UP | UV response: upregulated genes | 154 | 66 | -1.652 | 0.024 | Down | TAP1, CNP, APOM, CDKN1C, CYP1A1, UROD, FKBP4, EPHX1, SLC25A4, SIGMAR1, SQSTM1, NPTXR, H2AX, CCNE1, SULT1A1, IL6, RFC4, GRPEL1, CDC34, E2F5, HYAL2, STIP1, PSMC3, STK25, AP2S1, SPR, HMOX1, KCNH2, CHKA, FEN1, WIZ, MGAT1, AQP3, MRPL23, FURIN, RRAD, BSG, ATP6V1F, CCND3, IGFBP2, MAPK8IP2, DGAT1, MMP14, GAL, KLHDC3, STARD3, PARP2, JUNB, BAK1, RXRB, RPN1, TST, DNAJB1, ALDOA, YKT6, CLTB, SLC6A12, ICAM1, SELENOW, HLA-F, POLR2H, CDK2, NR4A1, NXF1, CYB5R1, ACAA1 |
| M1 | E2F TARGETS | cell cycle progression: E2F targets | 193 | 69 | -1.671 | 0.024 | Down | TK1, CDC20, SHMT1, MCM2, DDX39A, CDK4, KIF2C, HMGA1, MCM4, RANBP1, CKS1B, KIF22, RRM2, HMMR, BIRC5, TUBB, TRIP13, RNASEH2A, MCM3, MCM7, CCNB2, CDCA3, CDKN2C, CDKN2A, SPC24, PTTG1, H2AX, CCNE1, TOP2A, TIMELESS, CHEK1, SPAG5, DCTPP1, UBE2S, CENPM, CDK1, ESPL1, NAA38, ASF1B, DSCC1, TCF19, RPA2, SSRP1, LIG1, NASP, RFC2, KPNA2, WDR90, DLGAP5, UBE2T, MCM5, DNMT1, CDC25B, POP7, MYC, POLD1, MYBL2, ILF3, CIT, SNRPB, PSMC3IP, TBRG4, MKI67, TP53, PAICS, POLE, CDC25A, CDKN3, CENPE |
| M1 | DNA REPAIR | DNA repair | 143 | 65 | -1.772 | 0.024 | Down | POLR2E, ERCC2, NELFCD, ADA, TAF10, IMPDH2, TAF6, ZWINT, RAD52, ERCC1, AAAS, POLR1H, TARBP2, TYMS, NT5C, POM121, NELFE, NME4, RFC4, BRF2, SURF1, MRPL40, XPC, VPS37D, DDB2, RAD51, POLR2I, SUPT4H1, SUPT5H, VPS28, RPA2, SSRP1, GPX4, ALYREF, LIG1, POLD4, ITPA, RFC2, NELFB, SEC61A1, FEN1, NME3, GUK1, DGCR8, POLD1, ZNF707, ADRM1, GTF3C5, CLP1, ADCY6, MPG, EDF1, GTF2F1, TP53, RFC5, SAC3D1, DDB1, POLL, POLA2, APRT, NME1, PDE6G, PNP, CMPK2, POLR2J |
| M1 | MYC TARGETS V2 | "MYC targets, variant 2" | 57 | 35 | -1.882 | 0.024 | Down | CDK4, MCM4, WDR74, AIMP2, TCOF1, SLC19A1, DCTPP1, EXOSC5, TMEM97, MRTO4, PES1, NOP2, PHB1, GRWD1, RRP12, MYBBP1A, IMP4, MCM5, MAP3K6, MYC, RABEPK, SLC29A2, RRP9, DUSP2, TBRG4, PUS1, NOC4L, SRM, FARSA, NOP56, PA2G4, BYSL, RCL1, PLK4, GNL3 |
| M2 | INTERFERON GAMMA RESPONSE | interferon gamma response | 199 | 70 | -1.509 | 0.047 | Down | WARS1, BPGM, PTPN1, ICAM1, HLA-DMA, PSMB2, ADAR, IFITM2, MYD88, PNP, CASP7, TRIM21, CSF2RB, FPR1, NFKBIA, CDKN1A, PIM1, TXNIP, LAP3, PSME1, HLA-A, STAT2, STAT3, IDO1, TMT1B, IRF1, SERPING1, TRAFD1, IRF8, PTPN6, CD74, CMKLR1, GCH1, TNFAIP2, IRF2, NUP93, CCL5, RNF213, RNF31, LATS2, ARL4A, MX2, TRIM26, IL4R, PARP14, CMTR1, HLA-B, HLA-G, RIPK1, HLA-DRB1, IRF9, ZNFX1, NFKB1, RBCK1, PSMB9, NLRC5, CD86, UBE2L6, CD40, IFI35, PFKP, IL10RA, OGFR, MT2A, SLAMF7, ZBP1, SELP, IFITM3, HELZ2, MVP |
| M2 | COMPLEMENT | complement cascade | 194 | 56 | -1.548 | 0.036 | Down | FCER1G, PSEN1, PLAUR, GRB2, CTSB, RNF4, RABIF, RBSN, LGALS3, PIK3CG, CPQ, CDK5R1, CASP7, C4BPB, CDA, PRKCD, ITGAM, ANXA5, DUSP5, EHD1, C3, PRCP, FCN1, PIM1, GNAI2, LAP3, CTSS, PFN1, PLEK, IRF1, CTSD, PPP4C, RHOG, GNB2, SERPINA1, SERPING1, CASP9, GP9, CALM3, CD59, MSRB1, BRPF3, PIK3R5, LRP1, S100A12, CTSC, IRF2, SRC, LYN, TIMP2, CCL5, CA2, STX4, F5, CSRP1, XPNPEP1 |
| M2 | APOPTOSIS | programmed cell death; caspase pathway | 154 | 63 | -1.566 | 0.028 | Down | PAK1, DAP, GSR, PSEN1, PPP2R5B, IL1B, CTH, SQSTM1, H1-0, BCL2L1, LGALS3, BMF, IGF2R, SLC20A1, ETF1, GSN, CASP7, GSTM1, GPX4, GPX1, CDKN1A, BTG2, TXNIP, APP, IRF1, ADD1, DPYD, CASP9, CTNNB1, PLCB2, F2R, BNIP3L, MADD, GCH1, RNASEL, TIMP2, CASP2, JUN, EREG, TSPO, GUCY2D, DFFA, HMOX1, DAP3, AIFM3, LMNA, NEDD9, PDGFRB, RARA, CREBBP, CCND1, IFITM3, BAX, GADD45B, ATF3, IER3, RHOB, SPTAN1, GADD45A, ISG20, CD14, BIK, VDAC2 |
| M2 | PI3K AKT MTOR SIGNALING | PI3K signaling via AKT to mTORC1 | 103 | 35 | -1.643 | 0.025 | Down | AP2M1, ARF1, PRKAR2A, GRB2, CXCR4, SQSTM1, MAP2K3, MAPK1, DUSP3, TIAM1, MYD88, CALR, RPS6KA1, PRKCB, CDKN1A, NFKBIB, CFL1, SLC2A1, STAT2, SLA, PFN1, MKNK2, TNFRSF1A, PPP1CA, ECSIT, TRIB3, RALB, AKT1, RAC1, RIPK1, TSC2, AKT1S1, GRK2, YWHAB, SFN |
| M2 | IL6 JAK STAT3 SIGNALING | IL6 STAT3 signaling during acute phase response | 81 | 29 | -1.746 | 0.013 | Down | PTPN1, GRB2, IL1B, IFNGR2, CD9, CCR1, TNFRSF1B, MYD88, CSF2RB, IL17RA, PIM1, STAT2, STAT3, MAP3K8, IRF1, TNFRSF1A, ACVR1B, LEPR, PIK3R5, EBI3, TYK2, IL4R, JUN, CBL, IL10RB, ITGB3, IRF9, HMOX1, PF4 |
| M2 | REACTIVE OXYGEN SPECIES PATHWAY | reactive oxygen species pathway | 46 | 19 | -2.163 | < 0.001 | Down | PRDX2, CAT, PTPA, GSR, TXNRD2, PRDX6, PDLIM1, LSP1, OXSR1, MBP, PRDX1, GPX4, CDKN2D, MPO, NDUFS2, FTL, NDUFB4, LAMTOR5, FES |
| M2 | HEME METABOLISM | heme metabolism and erythroid lineage | 187 | 109 | -3.229 | < 0.001 | Down | FECH, ADD2, PRDX2, CAT, KEL, MFHAS1, DCAF11, SELENBP1, BPGM, HDGF, GYPB, AHSP, TAL1, PPP2R5B, OSBP2, TRIM10, E2F2, SLC4A1, SLC25A38, HAGH, CTSB, RAD23A, UROD, RCL1, FOXO3, SMOX, ADIPOR1, MAP2K3, RANBP10, RBM38, MARCHF2, UBAC1, H1-0, NCOA4, TENT5C, CA1, ASNS, EPB41, BLVRB, ISCA1, RNF123, BSG, SNCA, SLC25A37, GDE1, SEC14L1, SLC66A2, MARK3, ALAD, TRIM58, ANK1, MKRN1, TFDP2, EPB42, HMBS, ACP5, C3, RIOK3, NFE2, KAT2B, BTG2, MYL4, RAP1GAP, TNS1, DMTN, GYPC, KLF1, SLC2A1, ARHGEF12, DCUN1D1, CCND3, MARCHF8, TMCC2, MXI1, TCEA1, UCP2, PDZK1IP1, TRAK2, ADD1, HTATIP2, TMEM9B, FBXO9, GAPVD1, NFE2L1, LRP10, BNIP3L, FBXO7, SLC6A9, FOXJ2, TSPAN5, NARF, ENDOD1, H4C3, EIF2AK1, CLCN3, CA2, RHD, CCDC28A, UROS, HTRA2, PC, TOP1, LMO2, ICAM4, CDC27, MBOAT2, KLF3, GYPA, HBD |

### **Table S6.** Comprehensive temporally dynamic miRNA–mRNA regulatory pairs in preeclampsia; 29 predicted interactions involving miRNAs and target mRNAs dysregulated at both M1 (10–18 weeks) and M2 (32–38 weeks).

| **miRNA** | **miRNA PE Literature Evidence** | **Target Gene** | **Target PE Literature Evidence** | **M1 Direction** | **M2 Direction** | **miRNA Placenta** | **Target Placenta** | **VGLL3/Plazyo Connection**  (PMID:  41953989) |
| --- | --- | --- | --- | --- | --- | --- | --- | --- |
| miR-101-3p | Suppresses WDR5; reduces trophoblast proliferation/invasion (PMID: 36324256, 24804790, 32527308, 37437834) | ADGRA2 | Wnt7 co-receptor in trophoblasts/endothelium; decreased in early-onset PE; impairs proliferation/invasion, promotes apoptosis via JNK/P38 MAPK (PMID: 27016783, 38764331, 37694534) | miR ↑ target ↓ | miR ↓ target ↑ | ✔✔ | −✔ | STRONG: Wnt7/angiogenesis/invasion pathways directly overlap with Plazyo trophoblast differentiation defects (Fig 2A-B), endothelial dysfunction (Fig 3), PI3K/AKT dysregulation (Fig 4G); pathway-level overlap with Hippo/VGLL3-associated trophoblast or vascular programs |
| miR-15a-5p | Placental exosomal transfer promotes PE via PI3K/AKT/CDK1 (PMID: 33197745, 27529341) | ADGRA2 | Same as above | miR ↑ target ↓ | miR ↓ target ↑ | ✔✘ | −✔ | STRONG: PI3K/AKT pathway directly overlaps with Plazyo mTOR/PI3K findings (Fig 4G, S8D-E); exosomal transfer provides direct placental-to-blood communication link |
| miR-21-5p | Overexpressed in PE; detected in placenta and offspring (PMID: 26859593, 28694210, 33800458, 34611295, 30717412) | ADGRA2 | Same as above | miR ↑ target ↓ | miR ↓ target ↑ | ✔✔ | −✔ | STRONG: Multi-timepoint persistence; both miR-21-5p and ADGRA2 expressed in placenta; angiogenesis/invasion regulation connects to Plazyo endothelial dysfunction (Fig 3), trophoblast defects (Fig 2) |
| miR-26b-5p | Downregulated in severe PE placentas; reduced VDR, increased COX-2 (PMID: 34045549, 23830491, 19642860) | APOA2 | Indirect: HDL/lipid metabolism; limited direct PE evidence (PMID: 33674281 for GDM; APOA1: 38003549, 29914672, 37107170) | miR ↑ target ↓ | miR ↓ target ↑ | ✔✔ | −✔ | MODERATE: Metabolic dysregulation connects to Plazyo pathway findings |
| miR-26b-5p | Same as above | CEMP1 | Cementum protein; no direct PE association (dental/bone studies) | miR ↑ target ↓ | miR ↓ target ↑ | ✔✔ | ✔(Low)− | No clear Plazyo connection |
| miR-101-3p | Same as above | DEPP1 | 25261981 (DEPP1 as direct FOXO3 transcriptional target; ROS detoxification), 28545464 (DEPP1-mediated ROS accumulation as modulator of FOXO3-induced autophagy), 31125289 (dysregulated in PE), 29685721 (elevated in gestational diabetes) | miR ↑ target ↓ | miR ↓ target ↑ | ✔✔ | ✔(Low)✔ | DIRECT: DEPP1 is FOXO3-induced autophagy modulator; connects FOXO dysregulation (Plazyo Fig Fig 2D, 4G, refs 13-15), oxidative stress, and autophagy pathways. EXPERIMENTALLY VALIDATED. |
| miR-454-3p | Dysregulated in PE trophoblasts (PMID: 30367833, 30138897, 33129036) | DEPP1 | Same as above | miR ↑ target ↓ | miR ↓ target ↑ | ✔✔ | ✔(Low)✔ | Same FOXO-autophagy connection as above |
| miR-101-3p | Same as above | EEF2KMT | Elongation factor 2 kinase methyltransferase; protein synthesis regulation; no direct PE studies | miR ↑ target ↓ | miR ↓ target ↑ | ✔✔ | −✔(Low) | MODERATE: Translation regulation; may relate to Plazyo protein secretion pathway findings |
| miR-15a-5p | Same as above | EEF2KMT | Same as above | miR ↑ target ↓ | miR ↓ target ↑ | ✔✘ | −✔(Low) | Same as above |
| miR-106b-5p | Dysregulated in PE (PMID: 37184696, 35015290, 28545271) | EEF2KMT | Same as above | miR ↑ target ↓ | miR ↓ target ↑ | ✘✔ | −✔(Low) | Same as above |
| miR-19b-3p | PE trophoblast and placental dysregulation (PMID: 37215052, 32738709) | EEF2KMT | Same as above | miR ↑ target ↓ | miR ↓ target ↑ | ✔✘ | −✔(Low) | Same as above |
| miR-3178 | No direct PE literature; limited characterization | HSD17B1 | Catalyzes E1→E2 conversion; reduced in PE placentas; posttranscriptionally regulated by miR-210/518c; central dysregulated node in early-onset PE via scRNA-seq (PMID: 40394177, 22203747, 22786566, 38780059) | miR ↑ target ↓ | miR ↑ target ↑** | ✘✘ | ✔(High)✔ | EXCEPTION: Non-canonical regulation (both elevated M2); suggests alternative mechanisms. Estrogen biosynthesis critical for trophoblast differentiation connects to Plazyo differentiation defects |
| miR-106b-5p | Same as above | HSD17B1 | Same as above | miR ↑ target ↓ | miR ↓ target ↑ | ✘✔ | ✔(High)✔ | STRONG: Central dysregulated node in early-onset PE; estrogen synthesis essential for trophoblast differentiation overlaps with Plazyo differentiation defects (Fig 2A-B) |
| miR-20a-5p | PE placenta and trophoblast dysregulation (PMID: 22438230, 35870270, 26992682, 25210495, 41034780, 34963440) | HSD17B1 | Same as above | miR ↑ target ↓ | miR ↓ target ↑ | ✔✘ | ✔(High)✔ | Same as above |
| miR-15a-5p | Same as above | KCP | Kielin/chordin-like protein; suppresses TGF-β/activin via Smad2/3 (PMID: 16738323) | miR ↑ target ↓ | miR ↓ target ↑ | ✔✘ | −− | STRONG: TGF-β pathway regulation; Plazyo identifies TGFBR2 (ρ≥0.88 with VGLL3) and LTBP4 (TGF-β modulator) as VGLL3 targets (Fig 6G); TGF-β pathway enriched |
| miR-15a-5p | Same as above | NRM | Nurim (nuclear envelope membrane protein); no direct PE or placental studies | miR ↑ target ↓ | miR ↓ target ↑ | ✔✘ | −✔(Low) | No clear Plazyo connection |
| miR-20a-5p | Same as above | NRM | Same as above | miR ↑ target ↓ | miR ↓ target ↑ | ✔✘ | −✔(Low) | No clear Plazyo connection |
| miR-106b-5p | Same as above | NRM | Same as above | miR ↑ target ↓ | miR ↓ target ↑ | ✘✔ | −✔(Low) | No clear Plazyo connection |
| miR-26b-5p | Same as above | NRM | Same as above | miR ↑ target ↓ | miR ↓ target ↑ | ✔✔ | −✔(Low) | No clear Plazyo connection |
| miR-454-3p | Same as above | NRM | Same as above | miR ↑ target ↓ | miR ↓ target ↑ | ✘✘ | −✔(Low) | No clear Plazyo connection |
| miR-15a-5p | Same as above | NUP210L | Nuclear pore complex protein; no direct PE studies | miR ↑ target ↓ | miR ↓ target ↑ | ✔✘ | −✔(Low) | No clear Plazyo connection |
| miR-19b-3p | Same as above | NUP210L | Same as above | miR ↑ target ↓ | miR ↓ target ↑ | ✔✘ | −✔(Low) | No clear Plazyo connection |
| miR-106b-5p | Same as above | PHKG1 | Phosphorylase kinase gamma 1; glycogen metabolism; no direct PE studies | miR ↑ target ↓ | miR ↓ target ↑ | ✘✔ | −✔(Low) | MODERATE: Metabolic regulation; indirect connection to Plazyo metabolic findings |
| miR-19b-3p | Same as above | PHKG1 | Same as above | miR ↑ target ↓ | miR ↓ target ↑ | ✔✘ | −✔(Low) | Same as above |
| miR-20a-5p | Same as above | TLCD1 | TLC domain containing 1; ceramide synthesis; no direct PE studies | miR ↑ target ↓ | miR ↓ target ↑ | ✔✘ | ✔(High)− | MODERATE: Lipid metabolism; may relate to metabolic dysregulation |
| miR-106b-5p | Same as above | TLCD1 | Same as above | miR ↑ target ↓ | miR ↓ target ↑ | ✘✔ | ✔(High)− | Same as above |
| miR-15a-5p | Same as above | TSPAN16 | Tetraspanin 16; membrane protein; no direct PE studies | miR ↑ target ↓ | miR ↓ target ↑ | ✔✘ | −✔(Low) | No clear Plazyo connection |
| miR-101-3p | Same as above | UHRF1 | Ubiquitin-like PHD and RING finger domains 1; DNA methylation; epigenetic regulation (PMID: 35296332) | miR ↑ target ↓ | miR ↓ target ↑ | ✔✔ | ✔(Medium)− | MODERATE: Epigenetic regulation; may relate to Plazyo developmental trajectory findings |
| miR-15a-5p | Same as above | UHRF1 | Same as above | miR ↑ target ↓ | miR ↓ target ↑ | ✔✘ | ✔(Medium)− | Same as above |

**Footnotes:** miRNA–mRNA Regulatory Pairs: This table summarizes 29 predicted interactions between differentially expressed miRNAs and target mRNAs showing temporally overlapping dysregulation at both M1 (early pregnancy, 10–18 weeks) and M2 (late pregnancy, 32–38 weeks) in preeclamptic maternal whole blood compared to normotensive controls. Figure S1 includes selected directionally reversing protein-coding genes evaluated during network prioritization. Some genes shown in Figure S1, including C1orf54, were not retained in the final high-confidence miRNA–mRNA pair table because they did not meet target-prediction confidence criteria.

**Literature Evidence:** PMIDs (PubMed identifiers) support either miRNA or target gene association with preeclampsia, placental expression, or relevant functional mechanisms. 'Same as above' indicates reference to the first occurrence of that miRNA in the table to avoid redundancy.

Direction Notation: M1 and M2 columns show miRNA and target regulation patterns. 'miR ↑ / target ↓' indicates miRNA upregulated and target downregulated (canonical miRNA-mediated repression). 'miR ↓ / target ↑' indicates temporal reversal. **Exception: miR-3178 → HSD17B1 shows 'miR ↑ / target ↑**' at M2, representing non-canonical regulation suggesting alternative mechanisms (translational derepression, ceRNA sequestration, or post-translational stabilization).

**Placental Expression Notation:** First symbol = miTED database (for miRNA) or HPA database (for mRNA); second symbol = VDAART cohort (for both). ✔ = detected/confirmed; ✘ = not detected; − = no data available. Expression levels (High/Medium/Low) in parentheses indicate relative abundance when available from source databases. Genes marked ✔(Low) in the VDAART position were detected at low expression in VDAART placental RNA-seq.

**Source Databases:** miTED = miRNA Tissue Expression Database; VDAART = Vitamin D Antenatal Asthma Reduction Trial placental cohort; HPA = Human Protein Atlas.

**VGLL3/Plazyo Connection:** Contextual links to Plazyo et al. (Circulation 2026; PMID: 40502186), which implicated placental VGLL3, a Hippo pathway transcriptional coregulator, in preeclampsia-associated immune activation, trophoblast dysfunction, and endothelial pathology. Evidence strength: DIRECT = explicit mechanistic link with experimental support; STRONG = multiple pathway-level convergence with published data; MODERATE = pathway-level overlap; EXCEPTION = non-canonical regulatory mechanism. These annotations indicate pathway-level convergence with Plazyo/VGLL3 placental findings and should be interpreted as biological context rather than evidence that circulating signatures are directly caused by VGLL3-driven placental pathology.

Key Findings: (1) ADGRA2 targeted by 3 miRNAs (miR-101-3p, miR-15a-5p, miR-21-5p) with STRONG Plazyo connections via angiogenesis/trophoblast/PI3K pathways; (2) DEPP1 provides DIRECT connection to FOXO-autophagy axis dysregulated in Plazyo study and was experimentally validated; (3) HSD17B1 targeted by 3 miRNAs as central dysregulated node in early-onset PE; (4) KCP links to TGF-β pathway with direct VGLL3 target overlap (TGFBR2, LTBP4).

**Table S7.** Estimated leukocyte cell fractions by PE status and timepoint; No significant differences in major leukocyte populations (Bonferroni threshold: P<0.0023).

| **Cell Type**  (mean percentage ±SD) | **M1** | | **P Value** | **M2** | | **P Value** |
| --- | --- | --- | --- | --- | --- | --- |
|  | **PE Case**  **N = 41** | **Healthy**  **N = 82** |  | **PE Case**  **N = 38** | **Healthy**  **N = 97** |  |
| B Cell Memory | 0.53±1.02 | 0.46±0.77 | 0.660 | 0.62±1.04 | 0.59±1.06 | 0.892 |
| B Cell Naïve | 2.55±1.64 | 3.15±2.30 | 0.142 | 1.92±1.35 | 2.06±1.60 | 0.638 |
| B Cell Plasma | 0.14±0.23 | 0.19±0.60 | 0.575 | 0.20±0.31 | 0.31±0.57 | 0.246 |
| T Cell CD8+ | 6.07±4.59 | 6.84±4.44 | 0.369 | 6.53±5.63 | 5.67±4.48 | 0.356 |
| T Cell CD4+ Naive | 9.17±3.46 | 9.40±3.23 | 0.716 | 8.90±3.52 | 8.88±2.90 | 0.981 |
| T Cell CD4+ Memory Resting | 4.33±5.96 | 3.34±3.43 | 0.243 | 4.64±4.65 | 3.66±5.25 | 0.316 |
| T Cell CD4+ Memory Activated | 2.60±1.88 | 2.28±2.29 | 0.454 | 2.14±1.29 | 2.18±1.81 | 0.911 |
| T Cell Follicular Helper | 0 | 0 | NA | 0 | 0 | NA |
| T Cell Regulatory (Tregs) | 0.13±0.45 | 0.35±0.73 | 0.088 | 0.32±1.25 | 0.34±0.78 | 0.892 |
| T Cell Gamma Delta | 1.68±2.98 | 0.86±1.58 | 0.047 | 1.08±1.46 | 1.16±1.88 | 0.820 |
| NK Cell Resting | 1.48±1.53 | 1.98±2.11 | 0.176 | 1.69±1.74 | 1.95±2.07 | 0.494 |
| NK Cell Activated | 0.81±1.28 | 1.22±1.89 | 0.217 | 1.42±3.71 | 0.97±1.81 | 0.340 |
| Monocyte | 14.01±9.67 | 17.44±9.73 | 0.067 | 17.65±11.05 | 19.54±11.00 | 0.372 |
| Macrophage M0 | 0.04±0.23 | 0.24±0.79 | 0.111 | 0.10±0.37 | 0.39±1.06 | 0.104 |
| Macrophage M1 | 0 | 0 | NA | 0 | 0 | NA |
| Macrophage M2 | 1.39±1.59 | 1.07±1.38 | 0.243 | 1.26±1.43 | 0.97±1.31 | 0.269 |
| Myeloid Dendritic Cell Resting | 0.09±0.15 | 0.09±0.17 | 0.911 | 0.25±1.29 | 0.12±0.44 | 0.384 |
| Myeloid Dendritic Cell Activated | 0.92±0.66 | 0.86±0.69 | 0.671 | 0.89±0.62 | 0.90±0.66 | 0.967 |
| Mast Cell Resting | 0.68±1.12 | 0.65±0.98 | 0.883 | 0.73±1.04 | 0.74±1.01 | 0.994 |
| Mast Cell Activated | 1.21±1.59 | 0.90±1.35 | 0.261 | 1.19±1.54 | 0.91±1.27 | 0.290 |
| Eosinophil | 0 | 0 | NA | 0 | 0.03±0.27 | 0.522 |
| Neutrophil | 52.17±10.09 | 48.68±12.29 | 0.119 | 48.45±11.27 | 48.61±12.05 | 0.944 |

**Table S8.** Selection of miR-101-3p → DEPP1 for Functional Validation

| **Criterion** | **miR-101-3p** | **DEPP1** | **Evidence** |
| --- | --- | --- | --- |
| **Temporal dynamics** | M1: log₂FC=1.94, FDR=1.3×10⁻⁶ M2: Downregulated | M1: log₂FC=-2.32, FDR=0.0076 M2: Upregulated | Coordinated directional reversal |
| **Regulatory validation** | - | - | miRTarBase (experimental) TargetScan (computational) |
| **Placental expression** | Median log₂ = 13.1 (VDAART) IQR = 0.64 | Median log₂ = 10.76 (VDAART) IQR = 1.46 Vascular endothelial localization (HPA) | High expression, low variance |
| **Pathway connection** | - | FOXO1/FOXO3 target Enriched in 6 C8 endothelial signatures | Links to PI3K/AKT/mTOR reversal |
| **Biological function** | - | Autophagy modulator Oxidative stress response | Relevant to PE pathophysiology |
